# Interaction of PINK1 with nucleotides and kinetin

**DOI:** 10.1101/2023.08.08.552531

**Authors:** Zhong Yan Gan, Sylvie Callegari, Thanh N. Nguyen, Nicholas S Kirk, Andrew Leis, Michael Lazarou, Grant Dewson, David Komander

## Abstract

PINK1 is a ubiquitin kinase that accumulates on damaged mitochondria to trigger mitophagy, and PINK1 loss-of-function mutations cause early onset Parkinson’s disease. Nucleotide analogues such as kinetin triphosphate (KTP) have been suggested to enhance PINK1 activity and may represent a therapeutic strategy for the treatment of Parkinson’s disease. Here, we investigate the interaction of PINK1 with nucleotides, including KTP. We establish a cryo-EM platform exploiting the previously observed dodecamer assembly of *Pediculus humanus corporis* (*Ph*) PINK1 to determine PINK1 structures bound to AMP-PNP and ADP, which reveal unexpected conformational changes in the kinase N-lobe to enable PINK1 to form a ubiquitin binding site. Strikingly, we find that KTP is unable to bind *Ph*PINK1 or human (*Hs*) PINK1 due to a steric clash with the kinase ‘gatekeeper’ residue. Mutation of the gatekeeper to Ala or Gly is required to enable PINK1 to bind and utilise KTP as a phosphate donor in ubiquitin phosphorylation and mitophagy. Indeed, *Hs*PINK1 M318G can be used to conditionally uncouple PINK1 stabilisation and activity on mitochondria.

## Introduction

Parkinson’s disease (PD) is an incurable neurodegenerative disease, affecting more than 10 million individuals globally. While exact molecular mechanisms that cause the pathophysiology of PD remain unclear, much evidence suggests that a decline in mitochondrial health is a major contributor (*1*, *2*). The ubiquitin kinase PINK1 (encoded by *PARK6*/*PINK1*) and the E3 ubiquitin ligase Parkin (encoded by *PARK2*/*PRKN*) are amongst >15 *PARK*-encoded proteins that when mutated cause an early onset form of PD (EOPD), accounting for ∼5–10% of PD cases (*3–5*). PINK1 and Parkin are crucial mediators of mitophagy, a mitochondrial quality control pathway that degrades damaged mitochondria (*1*, *6*, *7*). PINK1 serves a key damage sensor and initiator of mitophagy. The kinase rapidly turns over under basal conditions (*8*), but upon mitochondrial depolarisation, accumulates on the outer mitochondrial membrane (OMM) where it forms a complex with the translocase of the outer membrane (TOM) and activates by autophosphorylation (*9–11*). Active PINK1 phosphorylates ubiquitin (*12–16*), enabling Parkin to be recruited to mitochondria by binding to phosphorylated ubiquitin (phospho-ubiquitin). PINK1 then contributes to activation of Parkin by phosphorylating its ubiquitin-like domain, leading to Parkin-mediated ubiquitination of OMM proteins and mitophagy (*1*, *6*).

Recent structural analysis of PINK1 from the body louse *Pediculus humanus corporis* (*Ph*) and the flour beetle *Tribolium castaneum* (*Tc*) provided detailed insights into the mechanism of PINK1 activation (*17–22*). PINK1 harbours a bilobal kinase fold, comprising N- and C-lobes that are embellished by three N-lobe insertions and helical extensions at its N- and C-termini (*17*, *19*). Structures of *Ph*PINK1 and *Tc*PINK1 dimers revealed that *trans*-autophosphorylation at a key Ser residue in PINK1 (Ser202/205/228 in *Ph*/*Tc*/*Hs*PINK1, respectively) activates the kinase (*17*, *19*). Furthermore, structures of phosphorylated *Ph*PINK1, with and without ubiquitin, demonstrated that autophosphorylation at the key Ser residue stabilises the third N-lobe insertion (insertion-3), enabling PINK1 to bind and phosphorylate ubiquitin (*17*, *18*). *Tc*PINK1 structures have been determined in complex with ATP analogues, which expectedly bind the cleft between the N- and C-lobes (*19*, *21*). However, structural changes as a consequence of nucleotide binding were not analysed or discussed.

Enhancing mitophagy by pharmacologically increasing PINK1 activity has been considered as a potential strategy to treat PD (*23*, *24*). In a key study, an analogue of ATP, kinetin triphosphate (KTP), was reported to be utilised by PINK1 with greater efficiency than ATP and restored the activity of a PINK1 EOPD mutant (*25*). While KTP itself is impermeable to the cell membrane and therefore of limited therapeutic value, its membrane permeable precursor, kinetin, can be intracellularly metabolised into KTP and appeared to accelerate PINK1/Parkin mitophagy (*25*, *26*). However, it remains unclear whether the PINK1 activating effect of kinetin is mediated through KTP, or whether an alternate mechanism is involved (*27*). Regardless, these compounds demonstrate considerable therapeutic potential, and it will be important to understand the molecular basis underlying their activities.

Here, we investigate PINK1’s interaction with the nucleotides AMP-PNP and ADP, and with the reported PINK1 activators, KTP and kinetin. We exploit a *Ph*PINK1 dodecamer as a platform to determine nucleotide-bound PINK1 structures by cryo-electron microscopy (cryo-EM). While structures of AMP-PNP-bound and ADP-bound *Ph*PINK1 reveal nucleotide-induced conformational changes in the PINK1 N-lobe, we failed to detect any binding between PINK1 and KTP. Instead, KTP binding is blocked by PINK1’s gatekeeper residue, which is analogous to many other kinases. Mutation of the Met gatekeeper residue to smaller residue enables KTP binding, and mutation to a Gly switches PINK1’s nucleotide preference from ATP to KTP, which inactivates PINK1 in cells. Mutated PINK1 can now be activated by treatment of cells with kinetin, and be used as a conditional activator of gatekeeper-mutated PINK1, and allows us to uncouple PINK1 stabilisation and PINK1 activity in mitophagy settings.

## Results

### Determining nucleotide-bound PINK1 structures by cryo-EM

We recently reported that a *Ph*PINK1 construct (residues 115–575) in its unphosphorylated state could assemble into a homo-dodecameric complex, enabling structural determination of the complex by cryo-EM (*17*). However, these structures of *Ph*PINK1 did not contain nucleotides, and were obtained with kinase inactive mutants or had undergone Cys-crosslinking procedures. While the arrangement of molecules within the dodecamer (six *Ph*PINK1 dimers) provided key insights into PINK1 activation, the *Ph*PINK1 dodecamer itself, with open and unobstructed ATP binding sites (Figure 1A), was the ideal platform to further determine whether and how nucleotide binding alters PINK1 conformation. We therefore sought to determine structures of wild-type (WT) *Ph*PINK1, without crosslinking, bound to ADP or the non-hydrolysable ATP analogue AMP-PNP.

**Figure 1.**
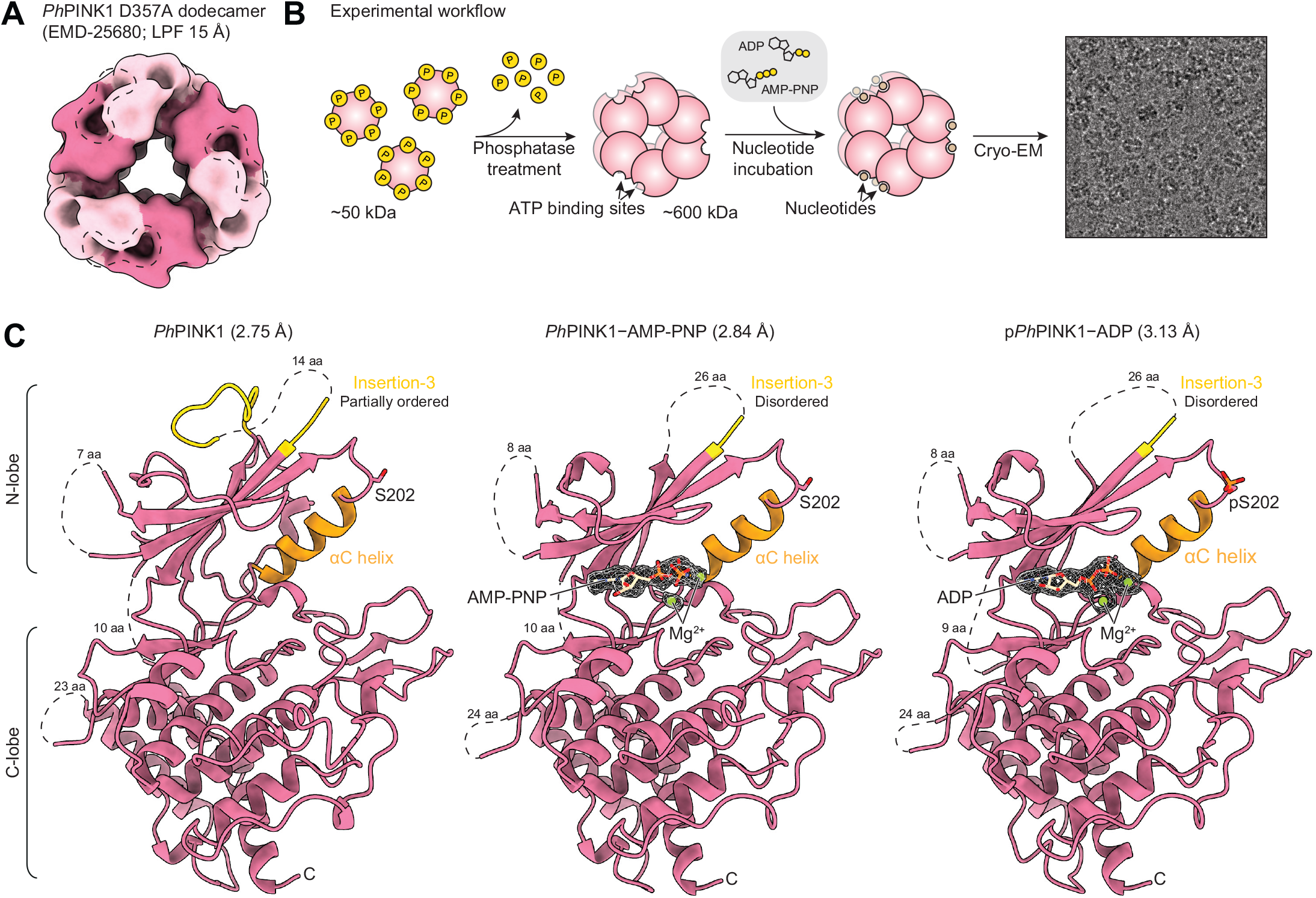
Determining nucleotide-bound *Ph*PINK1 structures. **(A)** 15-Å low-pass filtered (LPF) cryo-EM density map of the published *Ph*PINK1 D357A dodecamer ((*17*); EMDB-25680). Individual *Ph*PINK1 monomers are shown in alternating colours. Accessible ATP binding sites are highlighted in the dotted outlines (two binding sites per enclosed outline). **(B)** Workflow to generate the WT *Ph*PINK1 dodecamer for cryo-EM analysis in complex with nucleotides. The micrograph image is reused from Supplementary Figure 2C. **(C)** Structures of the nucleotide-free, AMP-PNP-bound and ADP-bound *Ph*PINK1 dimer (only chain B is shown, see Supplementary Figure 3 for whole dimers) at 2.75 Å, 2.84 Å and 3.13 Å resolution, respectively. Insertion-3 and the αC-helix are coloured in yellow and orange, respectively. Density of AMP-PNP, ADP and Mg^2+^ in the *Ph*PINK1 ATP binding site are shown as a mesh. aa, amino acids. Dotted lines indicate regions lacking electron density.

Thermal shift assays confirmed that ADP, ATP and AMP-PNP, each in the presence of Mg^2+^, bind and significantly stabilise monomeric *Ph*PINK1 (Supplementary Figure 1). To obtain WT *Ph*PINK1 dodecamers for cryo-EM analysis, *Ph*PINK1 was purified from bacteria in its monomeric and autophosphorylated form (*18*). Dephosphorylation using λ-phosphatase (λ-PP) induces oligomerisation into dodecamers that could be isolated and purified (Supplementary Figure 2A, see Materials and Methods). Phos-tag analysis, which resolves proteins according to phosphorylation status, confirmed that *Ph*PINK1 was homogeneously dephosphorylated (Supplementary Figure 2B), although a single phosphate remains on Thr305 due to the adjacent Pro306 that prevents dephosphorylation by λ-PP (Supplementary Figure 2B, 3C). The *Ph*PINK1 dodecamer was left untreated or was incubated with ADP/Mg^2+^ or AMP-PNP/Mg^2+^, prior to preparation of cryo-EM grids (Figure 1B, Supplementary Figure 2C, Materials and Methods). Processing in cryoSPARC (*28*) yielded reconstructions of the six-fold symmetric dodecamer, which was subsequently locally refined using a single dimer (asymmetric unit) to resolutions of 2.75 Å, 2.84 Å and 3.13 Å for the nucleotide-free, AMP-PNP-bound and ADP-bound *Ph*PINK1 dimers, respectively (Figure 1C, Supplementary Figure 2C, D, Table 1). Clear density could be observed for the bound nucleotides and Mg^2+^ ions (Figure 1C, Supplementary Figure 3A,B). The resolution of the maps enabled unambiguous modelling of *Ph*PINK1 with each nucleotide.

**Table 1.**
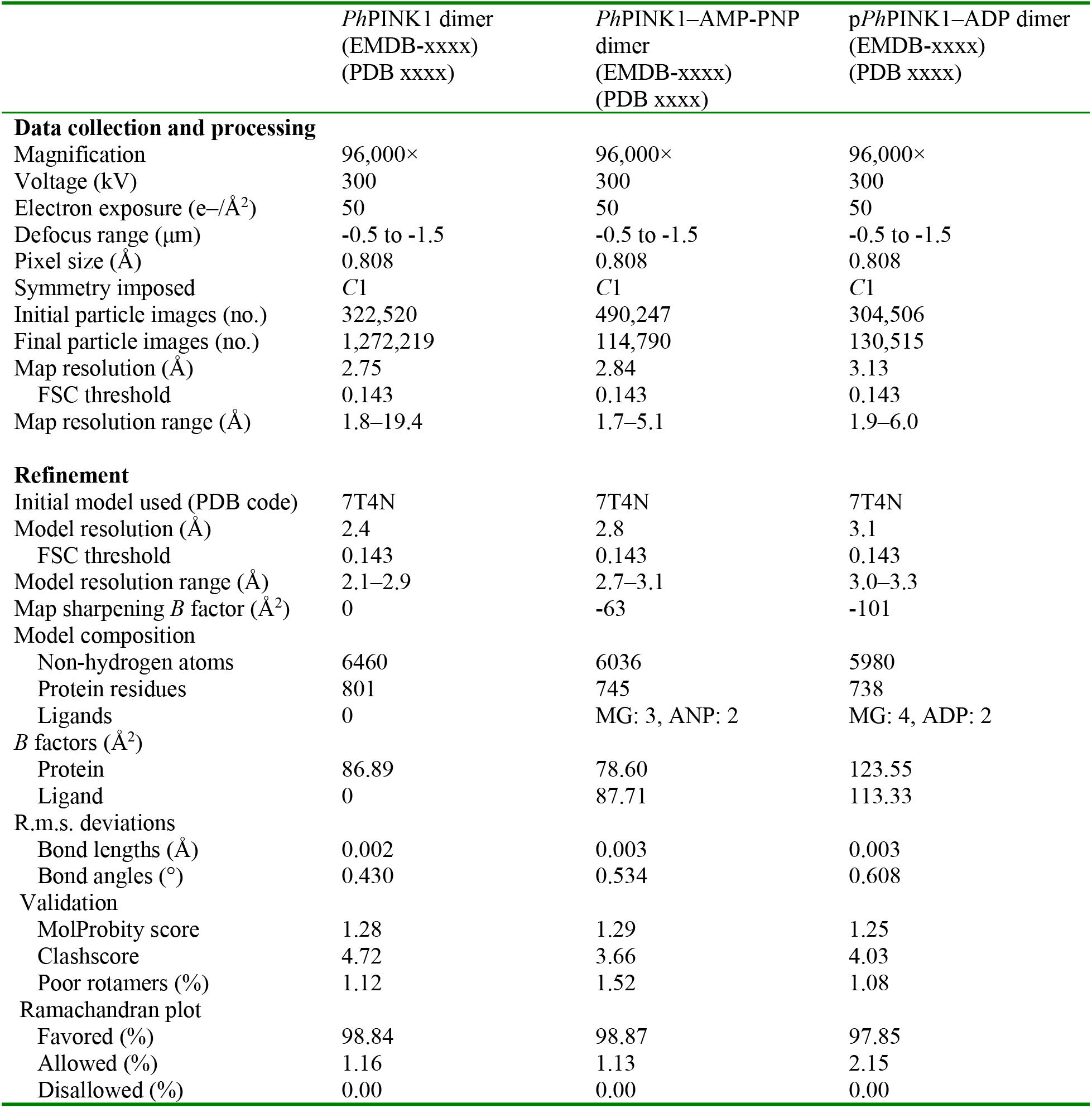
Cryo-EM data collection, refinement and validation statistics.

|  | *Ph*PINK1 dimer<br>(EMDB-xxxx)<br>(PDB xxxx) | *Ph*PINK1–AMP-PNP<br>dimer<br>(EMDB-xxxx)<br>(PDB xxxx) | p*Ph*PINK1–ADP dimer<br>(EMDB-xxxx)<br>(PDB xxxx) |
| --- | --- | --- | --- |
| **Data collection and processing** |  |  |  |
| Magnification | 96,000× | 96,000× | 96,000× |
| Voltage (kV) | 300 | 300 | 300 |
| Electron exposure (e<sup>-</sup>/Å<sup>2</sup>) | 50 | 50 | 50 |
| Defocus range (μm) | -0.5 to -1.5 | -0.5 to -1.5 | -0.5 to -1.5 |
| Pixel size (Å) | 0.808 | 0.808 | 0.808 |
| Symmetry imposed | C1 | C1 | C1 |
| Initial particle images (no.) | 322,520 | 490,247 | 304,506 |
| Final particle images (no.) | 1,272,219 | 114,790 | 130,515 |
| Map resolution (Å) | 2.75 | 2.84 | 3.13 |
| FSC threshold | 0.143 | 0.143 | 0.143 |
| Map resolution range (Å) | 1.8–19.4 | 1.7–5.1 | 1.9–6.0 |
| **Refinement** |  |  |  |
| Initial model used (PDB code) | 7T4N | 7T4N | 7T4N |
| Model resolution (Å) | 2.4 | 2.8 | 3.1 |
| FSC threshold | 0.143 | 0.143 | 0.143 |
| Model resolution range (Å) | 2.1–2.9 | 2.7–3.1 | 3.0–3.3 |
| Map sharpening *B* factor (Å<sup>2</sup>) | 0 | -63 | -101 |
| Model composition |  |  |  |
| Non-hydrogen atoms | 6460 | 6036 | 5980 |
| Protein residues | 801 | 745 | 738 |
| Ligands | 0 | MG: 3, ANP: 2 | MG: 4, ADP: 2 |
| *B* factors (Å<sup>2</sup>) |  |  |  |
| Protein | 86.89 | 78.60 | 123.55 |
| Ligand | 0 | 87.71 | 113.33 |
| R.m.s. deviations |  |  |  |
| Bond lengths (Å) | 0.002 | 0.003 | 0.003 |
| Bond angles (°) | 0.430 | 0.534 | 0.608 |
| Validation |  |  |  |
| MolProbity score | 1.28 | 1.29 | 1.25 |
| Clashscore | 4.72 | 3.66 | 4.03 |
| Poor rotamers (%) | 1.12 | 1.52 | 1.08 |
| Ramachandran plot |  |  |  |
| Favored (%) | 98.84 | 98.87 | 97.85 |
| Allowed (%) | 1.16 | 1.13 | 2.15 |
| Disallowed (%) | 0.00 | 0.00 | 0.00 |

### Nucleotide induced conformation changes in the PINK1 N-lobe

The newly generated *Ph*PINK1 structures revealed fresh insights into the interaction of nucleotides with PINK1, and unveiled significant conformational changes that couple nucleotide binding to previously observed conformational changes. The nucleotide-free WT *Ph*PINK1 dimer was virtually indistinguishable from the nucleotide-free *Ph*PINK1 D357A dimer we determined previously (*17*). Both AMP-PNP and ADP bind the ATP binding site of *Ph*PINK1 in the anticipated nucleotide binding mode, and in a similar manner to *Tc*PINK1 (Figure 2A, Supplementary Figure 3D) (*19*, *21*). Hydrophobic residues stemming from N- and C-lobes encapsulate the adenine and ribose groups (Figure 2A). Two hydrogen bonds are made via the N^1^ and N^6^ nitrogens of adenine, which contact the backbone of Tyr293 and Lys291, respectively, of the kinase hinge region (Figure 2A). The three phosphates of AMP-PNP extend towards the phosphoryl transfer centre of the kinase, and only indirectly bind the so-called P-loop (an extended Gly-rich β-hairpin loop including β1 and β2) of the N-lobe (Figure 2A). The phosphates are held in position by a series of electrostatic interactions with Lys193 of the kinase VAIK motif (LAVK in *Ph*PINK1), and two Mg^2+^ ions, which themselves are positioned by several polar residues that include Asp357 of the DFG motif (Figure 2A).

**Figure 2.**
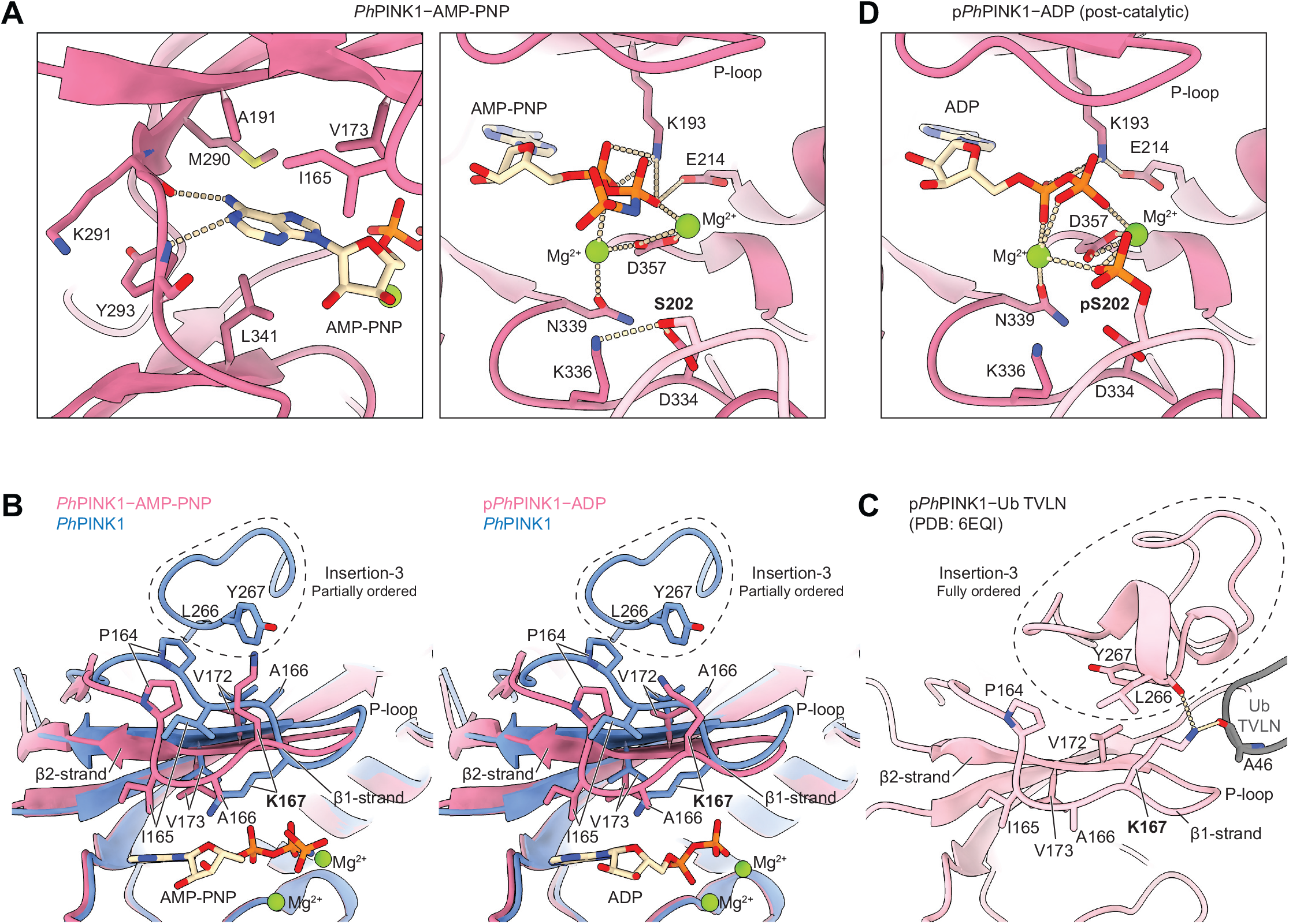
*Ph*PINK1-nucleotide interactions. **(A)** Details of the interaction between AMP-PNP in the ATP binding site of *Ph*PINK1 (chain B of the dimer). Interacting residues are shown as sticks, and polar interactions are shown as dotted lines. The interaction with Ser202 of chain A (light pink) is also shown. **(B)** Superimpositions of the N-lobe of AMP-PNP-bound and ADP-bound *Ph*PINK1 with nucleotide-free *Ph*PINK1, revealing differences in conformation of the P-loop and insertion-3 (highlighted in the dotted outlines). **(C)** The N-lobe of the published phosphorylated and ubiquitin-bound *Ph*PINK1 complex (Ub TVLN, ubiquitin T66V L67N mutant, PDB: 6EQI; (*18*)), in the same orientation as in **B**. **(D)** The interaction between ADP and the ATP binding site of *Ph*PINK1 (chain B of the dimer), and with phosphorylated Ser202 of chain A. The structure is displayed in the same orientation as the right panel of **A**.

Comparison of the nucleotide-free and nucleotide-bound states of *Ph*PINK1 revealed prominent reorganisation of the kinase P-loop upon nucleotide binding, which affects insertion-3 that eventually forms the ubiquitin binding site at the kinase N-lobe (Figure 2B). In nucleotide-free (and unphosphorylated) *Ph*PINK1, insertion-3 is only partially visible, but the ordered residues interact with, and shield, an otherwise exposed hydrophobic patch on the N-lobe, including Pro164, Ala166, Val172 of the P-loop (Figure 2B, Supplementary Figure 4A). This partially ordered conformation of insertion-3 is structurally incompatible with the fully ordered insertion- 3 formed after PINK1 N-lobe phosphorylation that becomes the binding site for ubiquitin and Ubl substrates (Figure 2C)(*17*, *18*). Interestingly, binding of AMP-PNP or ADP to unphosphorylated *Ph*PINK1 causes the P-loop to clamp onto the nucleotide. P-loop movement (by ∼6 Å between the Cα’s of Ile165) involves multiple residues, including Val173 of the β2- strand that forms part of the catalytic spine (C spine) and which is completed upon interaction with the nucleotide (Figure 2B) (*29*).

Two residues in the β1-strand, Ala166 and Lys167, flip such that Ala166 points its side chain into the ATP binding site, and Lys167 points towards the N-lobe (Figure 2B). As a net result, clamping down of the P-loop generates additional space for insertion-3 to fold, although without phosphorylation, insertion-3 remains disordered, and several hydrophobic side chains on the N-lobe, including the P-loop, are exposed (Figure 2B and Supplementary Figure 4A). Also important is the observed flip in Lys167, which only in the nucleotide bound or phosphorylated state of PINK1 can form interactions with both the ordered insertion-3, as well as with the substrate ubiquitin/Ubl (Figure 2C)(*17*, *18*).

Taken together, these observations indicate an unexpected and tight coupling between nucleotide binding status of PINK1 and the conformation of insertion-3 through conformational rearrangements of the kinase P-loop. Furthermore, given that phosphorylated *Ph*PINK1 in its nucleotide-free state adopts both active and inactive conformations (*17*), our structures suggest that ATP, in addition to its role as a phosphate donor, likely contributes to stabilising PINK1 in its active and ubiquitin binding-competent state (Supplementary Figure 5, see Discussion). The hydrophobicity of insertion-3 and residues in the P-loop are conserved in *Hs*PINK1, and we can easily envisage a similar mechanism in human PINK1 (Supplementary Figure 4B).

### A post-catalytic dimerised state of PINK1

To our surprise, we find that ADP-bound *Ph*PINK1 is autophosphorylated at Ser202, most likely due to contaminating ATP in the ADP stock we used (Figure 2D, Supplementary Figure 3E–G). This additional phosphate group interacts with both Mg^2+^ ions in the active site and is positioned ∼5 Å from the β-phosphate of the bound ADP (Figure 2D, Supplementary Figure 3G). The phosphorylated *Ph*PINK1–ADP dimer represents the post-catalytic state of PINK1 immediately following phosphoryl transfer, but prior to dimer dissociation. In this situation, *Ph*PINK1 molecules within the dimer remain entirely in their inactive conformation, with an extended αC- helix and disordered insertion-3, contrasting the conformational shift observed in our previous structure of phosphorylated and crosslinked *Ph*PINK1 dimer (*17*). A possible reason why the conformational change is not observed in this scenario could be that the high concentration of ADP in the sample sits snugly with phosphorylated pSer202 in the active site of a stable kinase dimer composition (Supplementary Figure 3G).

### PINK1 cannot use KTP due to a clash with a gatekeeper residue

We next attempted to investigate how the reported PINK1 activator kinetin triphosphate (KTP; Figure 3A) interacts with PINK1 (*25*). Using thermal shift binding assays, we first tested whether KTP stabilises *Ph*PINK1, and observed that unlike ATP, KTP does not stabilise *Ph*PINK1 (Figure 3B and Supplementary Figure 6C). Next, *in vitro* ubiquitin phosphorylation assays showed that *Ph*PINK1 could not phosphorylate ubiquitin when KTP was the sole nucleotide source (Figure 3C). Given that the previous study reporting PINK1 activity with KTP was based on human PINK1 (*25*), we tested whether sequence differences between *Ph*PINK1 and *Hs*PINK1 (∼40% kinase domain identity) may account for the inability of KTP to work with *Ph*PINK1. To minimise these differences, we mutated the three differing residues within the *Ph*PINK1 ATP binding site to *Hs*PINK1 equivalent residues (V247A, M249V, T356A), generating a humanised version of *Ph*PINK1 (Supplementary Figure 6A). Despite now harbouring a *Hs*PINK1-like ATP binding site, the *Ph*PINK1 V247A/M249V/T356A triple mutant was neither stabilised nor showed ubiquitin kinase activity with KTP (Supplementary Figure 6B–D).

**Figure 3.**
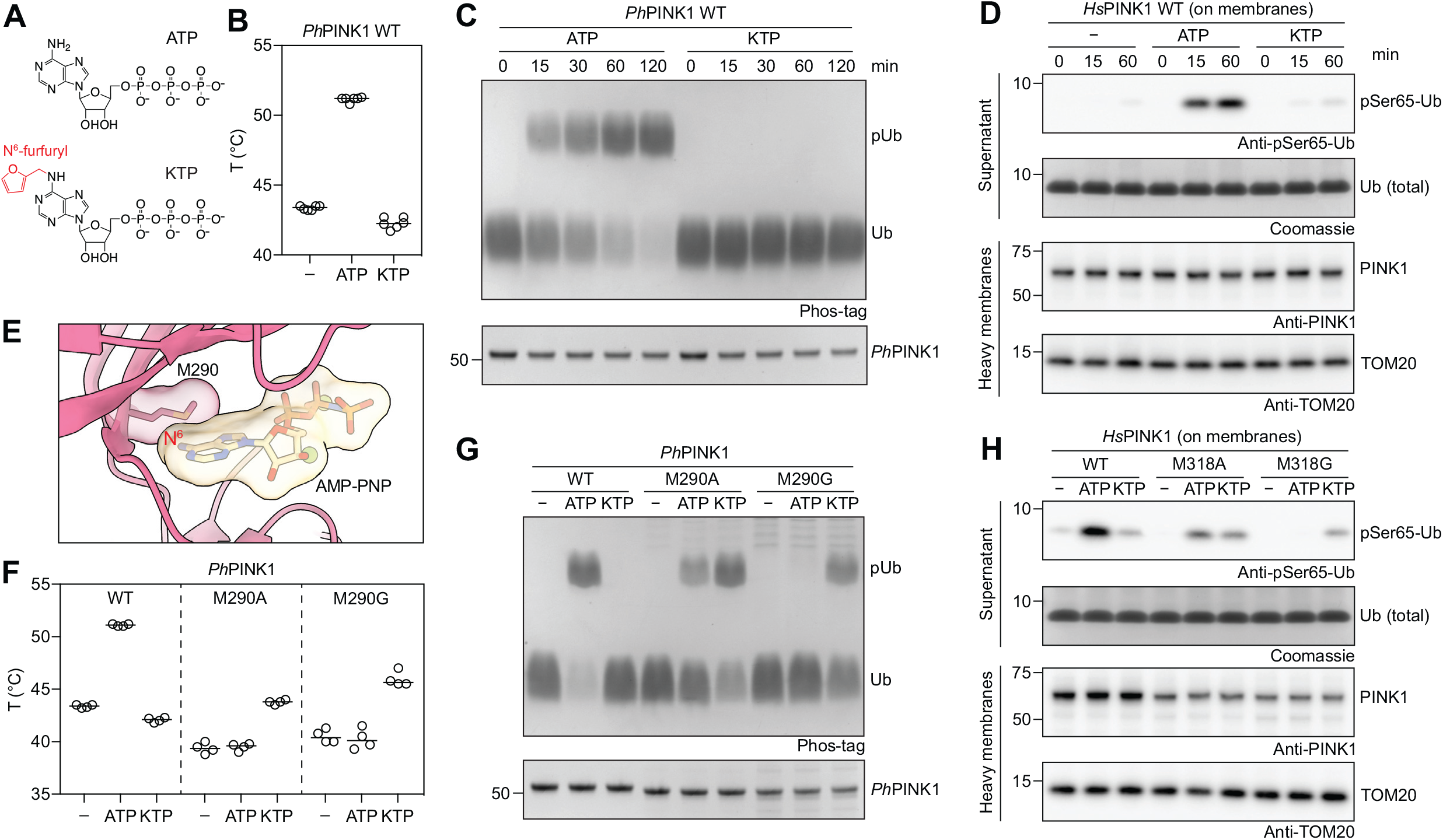
The PINK1 gatekeeper residue prevents KTP binding. **(A)** Chemical structures of ATP and KTP. The N^6^-furfuryl group of KTP is highlighted in red. **(B)** Melting temperatures of WT *Ph*PINK1 (residues 115–575) in the presence of ATP or KTP. Experiment was performed three times in technical duplicates. **(C)** Time course ubiquitin phosphorylation assay using WT *Ph*PINK1 in the presence of ATP or KTP and analysed by Phos-tag SDS-PAGE. Experiment was performed in triplicate. **(D)** Time course ubiquitin phosphorylation assay using *Hs*PINK1-containing heavy membranes from OA-treated HeLa cells in the presence of ATP or KTP (see Materials and Methods) and analysed by Western blotting. Experiment was performed in triplicate. **(E)** The ATP binding site of AMP-PNP-bound *Ph*PINK1, revealing that the gatekeeper Met290 would sterically block an N^6^-modified ATP analogue from binding *Ph*PINK1. **(F)** Melting temperatures of *Ph*PINK1 WT and gatekeeper mutants M290A and M290G in the presence of ATP or KTP. Experiment was performed two times in technical duplicates, while M290A in the absence of nucleotide was performed three times in technical duplicates (see Supplementary Figure 7). **(G)** Ubiquitin phosphorylation assay using *Ph*PINK1 WT and gatekeeper mutants M290A and M290G in the presence of ATP or KTP for 2 h and analysed by Phos-tag SDS-PAGE. Experiment was performed in triplicate. **(H)** Ubiquitin phosphorylation assay using *Hs*PINK1-containing heavy membranes from OA-treated HeLa cells in the presence of ATP or KTP for 2 h (see Materials and Methods) and analysed by Western blotting. Experiment was performed in triplicate.

Next, we investigated *Hs*PINK1 activity, using heavy membranes from OA-treated HeLa *PINK1*^−/−^ cells transiently expressing WT *Hs*PINK1 that were incubated with recombinant ubiquitin in the presence of either ATP or KTP (see Materials and Methods). Incubation with ATP resulted in phosphorylation of ubiquitin at Ser65, as detected by a phospho-Ser65 ubiquitin antibody (Figure 3D). While KTP incubation resulted in a faint phospho-ubiquitin band, a similarly faint band was also visible in the absence of any added nucleotide, indicating the presence of residual ATP in the crude heavy membrane preparation used for the assay (Figure 3D). These results indicate that PINK1 is unable to use KTP as a phosphate donor.

Why was PINK1 unable to use KTP in our experiments? KTP is an analogue of ATP, defined by an additional furfuryl group covalently attached to the N^6^ of the adenine ring (Figure 3A). N^6^- substituted ATP analogues with bulky groups such as furfuryl can typically not be accommodated by protein kinases due to a clash with a so-called gatekeeper residue at the back of the ATP binding site (*30*). It is possible that the gatekeeper residue of PINK1 (Met290 in *Ph*PINK1, Met318 in *Hs*PINK1) obstructs KTP (Figure 3E). Indeed, mutation of Met290 in *Ph*PINK1 to smaller Ala or Gly residues, while impacting recombinant protein yield and stability (Supplementary Figure 7), enabled the kinase to be stabilised by KTP, and to use KTP as a phosphate donor in ubiquitin phosphorylation experiments (Figure 3F, G). These results were mirrored in *Hs*PINK1 M318A and M318G mutants enriched from OA-treated HeLa cells (Figure 3H). While both kinases were expressed at lower levels (Figure 3H), the Gly mutation greatly diminished PINK1’s ability to utilise ATP, consistent with the idea that the gatekeeper is also important for ATP binding (Figure 3G, H). However, both mutants were able to use KTP instead of ATP to generate phospho-ubiquitin.

Taken together, our results show that contrary to what has been suggested (*25*, *31*, *32*) PINK1 may not bind KTP or use it as a preferred phosphate donor in a direct ATP-competitive fashion. Importantly, we could enable PINK1 to utilise KTP by mutating the gatekeeper Met residue in the PINK1 ATP binding site in insect and human PINK1. Switching PINK1’s nucleotide preference from ATP to KTP was next exploited to decouple PINK1 stabilisation from PINK1 activity.

### Kinetin activates human PINK1 gatekeeper mutants in cells

We assessed the impact of the gatekeeper mutants M318A and M318G on *Hs*PINK1 function in intact HeLa *PINK1*^−/−^ cells expressing YFP–Parkin that were transiently transfected with constructs encoding *Hs*PINK1 variants. *Hs*PINK1 WT accumulated in response to OA and generated phospho-ubiquitin, as expected (Figure 4A). Accumulation of PINK1 was also observed for *Hs*PINK1 M318A and M318G mutants, and proteasome inhibition induced similar accumulation of the 52-kDa PARL-cleaved PINK1 fragment, indicating that basal turnover of PINK1 variants was unimpaired (Figure 4A). While a *Hs*PINK1 M318A mutant phosphorylated ubiquitin at slightly reduced levels compared with WT, the M318G mutant generated phospho-ubiquitin at barely detectable levels (Figure 4A), consistent with *in vitro* experiments (Figure 3H).

**Figure 4.**
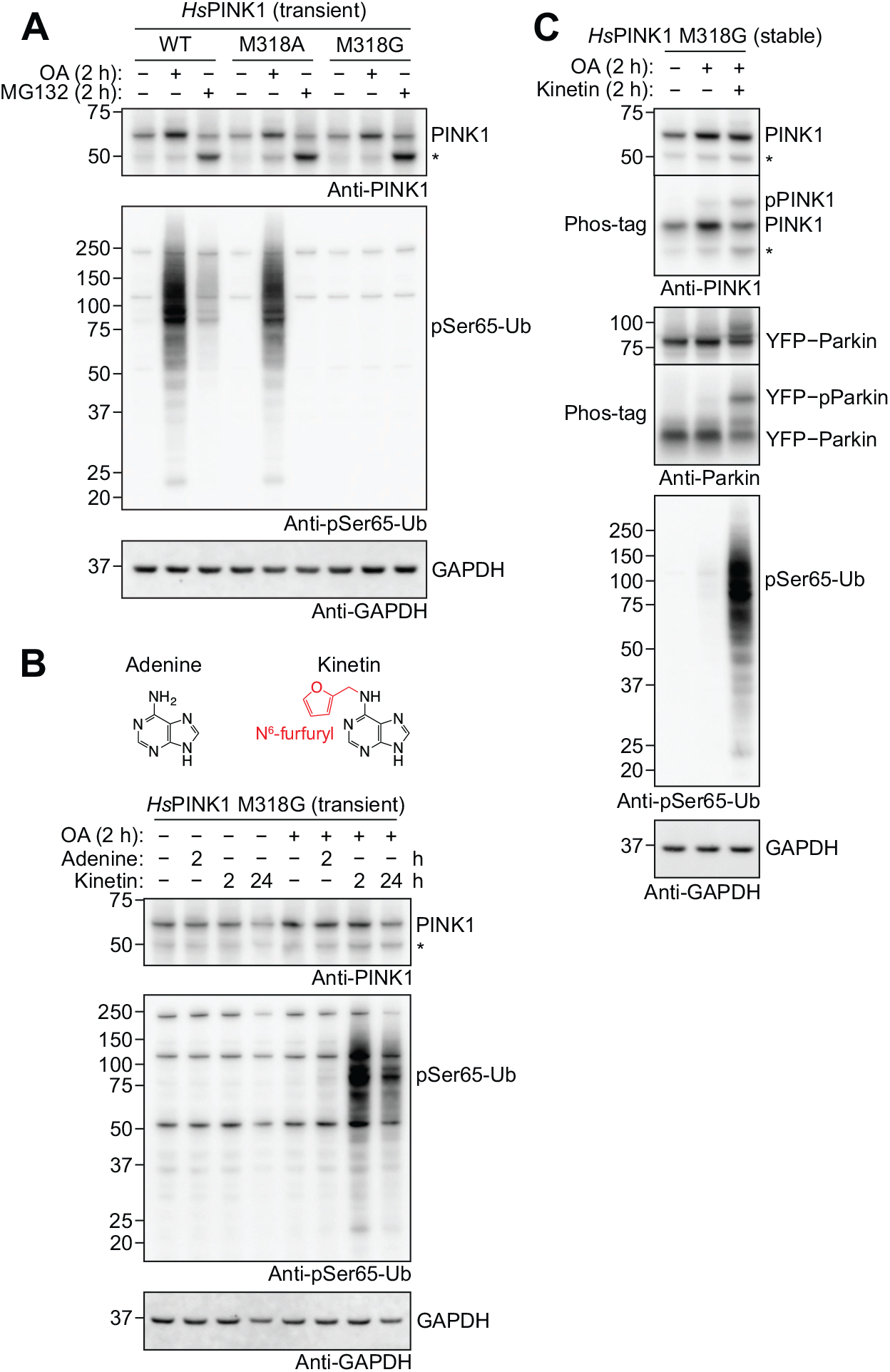
Kinetin activates *Hs*PINK1 M318G in cells. **(A)** HeLa *PINK1*^−/−^ YFP–Parkin cells transiently expressing *Hs*PINK1 WT, M318A and M318G were treated with OA or MG132 for 2 h, then immunoblotted for PINK1 and phospho-Ser65- ubiquitin (pSer65-Ub). *Hs*PINK1 M318G is stabilised upon OA treatment but is unable to generate phospho-ubiquitin. Experiment was performed in triplicate. **(B)** HeLa *PINK1*^−/−^ YFP– Parkin cells transiently expressing *Hs*PINK1 M318G were treated with 200 µM adenine for 2 h, or 200 µM kinetin for 2 h or 24 h. OA was added 2 h before lysis. Immunoblotting revealed that the addition of kinetin co-treatment with OA activates *Hs*PINK1 M318G and leads to ubiquitin phosphorylation. Experiment was performed in triplicate. **(C)** HeLa *PINK1*^−/−^ YFP–Parkin cells stably expressing *Hs*PINK1 M318G treated for 2 h with OA alone, or in the presence of 200 µM kinetin. As in **B**, immunoblotting revealed that kinetin activates ubiquitin phosphorylation. Additional Phos-tag analysis shows induction of PINK1 autophosphorylation and Parkin phosphorylation. Experiment was performed in triplicate. An asterisk in **A–C** indicates the 52-kDa PARL-cleaved PINK1.

Given the M318G mutant’s preference for KTP over ATP, we wondered whether its inactivity in cells can be overcome by supplying KTP. However, direct KTP treatment is unfeasible as ATP analogues are unable to cross the cell membrane. Instead, the KTP precursor, kinetin, is membrane permeable and has been shown to be intracellularly metabolised into KTP (*25*). We therefore attempted to activate *Hs*PINK1 M318G with kinetin. Treatment with OA alone for 2 h did not activate the M318G mutant, but co-treatment with 200 µM kinetin, but not adenine, led to substantial generation of phospho-ubiquitin (Figure 4B). Increasing the duration of kinetin treatment to 24 h did not increase the level of phospho-ubiquitin, but instead hampered cell growth and/or survival (Figure 4B). Kinetin did not enhance the activity of *Hs*PINK1 WT, suggesting the effect of kinetin is specific to *Hs*PINK1 M318G (Supplementary Figure 8). These results are consistent with kinetin undergoing intracellular conversion into KTP, which then acts as a phosphate donor specifically for the *Hs*PINK1 M318G gatekeeper mutant.

We also generated cells stably expressing *Hs*PINK1 M318G. As was observed in transient expression experiments, stably expressed *Hs*PINK1 M318G accumulated in response to OA treatment and generated barely detectable levels of phospho-ubiquitin unless co-treated with kinetin (Figure 4C). Phos-tag analysis further revealed that kinetin co-treatment significantly increased PINK1 autophosphorylation and Parkin phosphorylation (Figure 4C).

### Kinetin-activated PINK1 M318G recruits Parkin to mitochondria

Since Parkin must be in proximity to OMM-stabilised PINK1 to become phosphorylated, our results suggested that kinetin-activated *Hs*PINK1 M318G could induce translocation of Parkin to mitochondria and trigger mitochondrial clearance via mitophagy. To test this, we performed immunofluorescence using HeLa *PINK1*^−/−^ cells expressing YFP–Parkin and *Hs*PINK1 to assess YFP–Parkin translocation to mitochondria. As anticipated, by the 1 h timepoint of OA treatment, WT *Hs*PINK1 had robustly recruited YFP–Parkin to mitochondria (Figure 5). In contrast, *Hs*PINK1 M318G did not recruit Parkin after 1 h of OA treatment, however, when co-treated with kinetin, robust recruitment was observed at 1 h OA/kinetin treatment (Figure 5). Taken together, the results indicate that *Hs*PINK1 M318G is functionally compromised to induce mitophagy in normal cells, but now relies on a distinct nucleotide source, KTP, which is generated *in situ* from kinetin. We have hence established an orthogonal system to induce mitophagy, in which PINK1 stabilisation and PINK1 activation are uncoupled.

**Figure 5.**
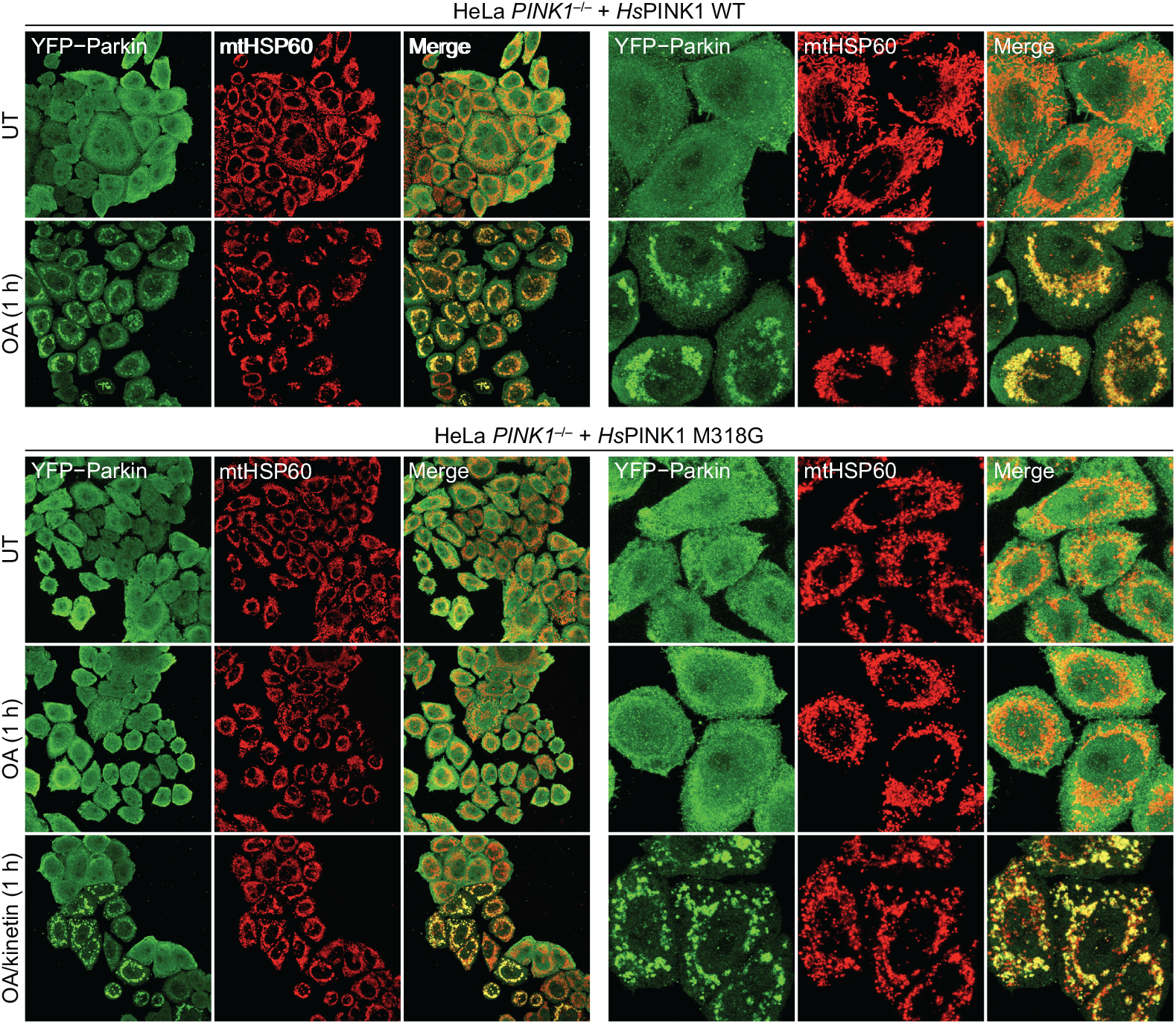
Kinetin-activated *Hs*PINK1 M318G induces Parkin translocation. YFP–Parkin translocation in fixed HeLa *PINK1*^−/−^ cells stably expressing *Hs*PINK1 WT or M318G and immunostained for mtHSP60 (mitochondrial marker). Cells were treated with either OA alone or in combination of 200 µM kinetin for 1 h prior to fixing and staining. While YFP– Parkin translocation was induced with OA alone in cells expressing *Hs*PINK1 WT, translocation in *Hs*PINK1 M318G-expressing cells was only induced when co-treated with kinetin. However, the M318G mutant is not completely inactive; when cells were treated with OA alone for 2 h, YFP–Parkin translocation was observable (see Supplementary Figure 9). Experiment was performed in duplicate with Supplementary Figure 9, and representative images are shown.

## Discussion

In the first part of this study, we exploited established cryo-EM workflows to characterise the interaction of *Ph*PINK1 with nucleotides. New structures reveal localised conformational changes that occur upon nucleotide binding (Figure 2). While crystal structures of nucleotide-free and AMP-PNP-bound *Tc*PINK1 have been reported previously (*19–21*), a direct comparison between nucleotide-free and nucleotide bound states was not performed. We used cryo-EM to directly compare the nucleotide-bound and unbound states of *Ph*PINK1, revealing conformational changes in the kinase P-loop and insertion-3 (Supplementary Figure 5). Prior to nucleotide binding, insertion-3 shields a hydrophobic patch in the N-lobe, likely to maintain protein stability and prevent non-specific interactions with other proteins prior to the activation of PINK1. Nucleotide binding is relayed via the flexible P-loop, and opens the N-lobe to accommodate insertion-3; however, without phosphorylation, insertion-3 remains disordered. Such exposed state of PINK1 would be short-lived since subsequent *trans*-autophosphorylation of PINK1 would cause insertion-3 to reconfigure into its activated and ubiquitin binding conformation, thereby re-shielding the hydrophobic patch (*17–19*). Furthermore, following autophosphorylation, insertion-3 remains dynamic and exchanges between folded and unfolded states, as we have previously resolved using 3D variability analysis (*17*). Therefore, it is likely that ATP binding, in addition to it serving as a phosphate donor, helps to further stabilise insertion-3 in its folded and ubiquitin binding conformation, increasing the efficiency of ubiquitin binding and phosphorylation.

The ease with which the nucleotide-bound *Ph*PINK1 structures could be solved means that the *Ph*PINK1 dodecamer could now be used as a platform to understand the binding mechanism of other kinase-interacting molecules, such as PINK1-activating compounds. We had envisaged to use our platform to understand how KTP interacts with PINK1, however we could not biochemically reproduce the reported PINK1-binding and activating properties of KTP (*25*). With our detailed structural understanding of PINK1, we define exactly why KTP cannot interact with PINK1: the PINK1 gatekeeper residue prevents the enlarged nucleobase to bind in the ATP binding pocket due to a steric clash between KTP furfuryl group and the gatekeeper Met side chain. Importantly, mutation of the gatekeeper to the smaller residue such as Ala and Gly enables KTP to bind and activate PINK1. These results are consistent with numerous reports of kinases that require a gatekeeper mutation to use KTP (in the KTPγS form) as their phosphate source (*33–38*). To the best of our knowledge, apart from PINK1, there have been no reports of kinases that are able to accommodate KTP in their unmodified WT form.

The KTP precursor kinetin and its derivatives have been thought to activate PINK1 in cells by undergoing intracellular conversion into KTP to function as a phosphate donor (*25*, *26*, *32*). Our data now indicates that the underlying mechanism behind kinetin induced PINK1 activation is unlikely to occur via KTP providing an improved nucleotide source to PINK1. A recent study reported that a kinetin analogue MTK458, which cannot be converted into a triphosphate form, was able to activate PINK1 activity (*27*). Our data would agree that MTK458 is unlikely to act as an ATP-analogue/phosphate donor in the PINK1 kinase reaction. How MTK458 acts on PINK1 will require further analysis and may also illuminate the role of kinetin in PINK1 biology.

Interestingly, kinetin or MTK458 appear to work more efficiently in combination with sub-threshold doses of depolarising agents that are insufficient to trigger phospho-ubiquitin generation and mitophagy on their own (*26*, *27*). We used kinetin in combination with typical mitophagy-inducing doses of the depolarising agent OA, but did not detect kinetin-mediated amplification of *Hs*PINK1 activity. Instead, we saw a remarkable sensitisation to kinetin-mediated activation when the M318G gatekeeper mutation was introduced. The specificity of kinetin toward the M318G mutant strongly indicates that the mechanism underlying PINK1 activation in our case is via the conversion of kinetin into KTP, which then acts directly on PINK1 M318G via its enlarged ATP binding site.

Since *Hs*PINK1 M318G accumulates on mitochondria upon depolarisation, yet remains almost completely inactive, kinetin may be used to specifically activate pre-accumulated *Hs*PINK1 M318G, essentially decoupling PINK1 activity from stabilisation of the protein through mitochondrial depolarisation. A similar goal has been achieved previously using a temperature sensitive mutant of PINK1 that becomes active when the temperature is lowered from 37 °C to 22 °C (*39*). Our strategy, while not needing a temperature shift, requires the initial conversion of kinetin into KTP. Based on our immunofluorescence imaging experiments, conversion to KTP produces an effect on Parkin recruitment within 1 h. If a more rapid conversion of kinetin is required, it may be possible to use kinetin riboside derivatives that reduce the number of conversion steps into KTP (*32*). We note that *Hs*PINK1 M318G is not a kinase inactive mutant since it can function with ATP when expressed at high levels and subjected to extended OA treatments. Therefore, *Hs*PINK1 M318G expression should be titrated to a level that induces minimal OA-induced Parkin recruitment and mitophagy, while maintaining robust activation of kinase activity by kinetin.

The second kinase gatekeeper mutant we used, *Hs*PINK1 M318A, is active with both ATP and KTP, and remains functional in the absence of kinetin. This mutant has been used previously for the purpose of sensitising PINK1 to inhibition by PP1 analogues such as 1-NA-PP1 and 1-NM-PP1 that were specifically designed specifically to inhibit gatekeeper-mutated kinases (*40*, *41*). It is likely that *Hs*PINK1 M318G may also be inhibited by PP1 analogues, given that PP1 analogues inhibit many kinases harbouring a Gly gatekeeper mutation (*40*). Therefore, kinetin and PP1 analogues, in combination with the *Hs*PINK1 M318A and M318G gatekeeper mutants, may comprise a powerful chemical genetics toolkit for the activation or inhibition of PINK1 activity in cells in an experimentally controlled manner.

## Materials and Methods

### Molecular cloning

DNA encoding *Ph*PINK1 (residues 115–575), codon optimised for expression in *Escherichia coli*, was inserted in between the KpnI and HindIII sites of the pOPINK vector (*42*) using the In-Fusion HD Cloning Kit (Takara). The pOPINK vector incorporates an N-terminal GST tag and a 3C protease cleavage site into the *Ph*PINK1 construct. *Ph*PINK1 mutants were generated using the Q5 Site-Directed Mutagenesis Kit (NEB).

### Protein expression and purification

All *Ph*PINK1 (residues 115–575) constructs and λ-PP were expressed in *Escherichia coli* Rosetta2 (DE3) pLacI cells (Novagen) and purified as described previously (*17*). To generate the WT *Ph*PINK1 dodecamer for cryo-EM analysis, ∼11 mg of purified WT *Ph*PINK1 (residues 115–575), at a concentration of 15 μM, was dephosphorylated with 7.5 μM λ-PP in 25 mM Tris (pH 8.5), 500 mM NaCl, 2 mM MnCl_2_, 5% (v/v) glycerol, 10 mM DTT for 24 h at 4 °C. To promote *Ph*PINK1 oligomerisation, the concentration of NaCl was reduced by buffer exchange into 25 mM Tris (pH 8.5), 150 mM NaCl, 10 mM DTT using a HiPrep 26/10 Desalting column (Cytiva), then incubated for 3 h at 4 °C. The resulting *Ph*PINK1 dodecamer was purified on a HiLoad 26/600 Superdex 200 pg column (Cytiva) in 25 mM Tris (pH 8.5), 150 mM NaCl, 10 mM DTT, and fractions corresponding to the dodecamer were pooled. Anion exchange chromatography was used to concentrate the dodecamer. Pooled fractions from SEC were applied to a Mono Q 5/50 GL column (Cytiva) in 25 mM Tris (pH 8.5), 50 mM NaCl, 10 mM DTT and eluted with a 0–50% linear gradient of 25 mM Tris (pH 8.5), 1 M NaCl, 10 mM DTT over 20 column volumes. The *Ph*PINK1 dodecamer eluted at approximately 250 mM NaCl. The fraction containing the highest concentration of *Ph*PINK1 (3.4 mg/mL) was diluted with 25 mM Tris (pH 8.5), 10 mM DTT to achieve a 150 mM NaCl concentration, resulting in a final *Ph*PINK1 concentration of 1.9 mg/mL. The protein was then flash-frozen in liquid nitrogen and stored at −80 °C.

### Cryo-EM sample preparation and data collection

To generate the *Ph*PINK1–nucleotide complexes, purified *Ph*PINK1 dodecamer (1.9 mg/mL) was incubated with 10 mM AMP-PNP or ADP and 10 mM MgCl_2_ for 10–25 min prior to vitrification. The nucleotide-free or nucleotide-bound dodecamers were dispensed onto glow discharged UltrAuFoil (Quantifoil GmbH, Germany) R1.2/1.3 holey specimen support (‘grid’) at 100% humidity, 4 °C, and blotted for 4 s (nominal blot force −1). Grids were then plunge frozen in liquefied ethane using a Vitrobot Mark IV (Thermo Fisher Scientific). Data were collected using a Titan Krios G4 microscope equipped with a Falcon 4 direct electron detector (Thermo Fisher Scientific) using a nominal magnification of 96,000×, corresponding to a pixel size at the detector of 0.808 Å. A total of 2,746, 3,094 and 2,923 movies were captured for the nucleotide-free, AMP-PNP-bound and ADP-bound *Ph*PINK1 datasets, respectively.

### Cryo-EM refinement and model building

Cryo-EM processing was performed in cryoSPARC (v4.2.1) (*28*). All *Ph*PINK1 datasets were processed using a similar workflow, detailed in Supplementary Figure 2C. All movies were patch-motion corrected, and CTF parameters were estimated using the patch CTF job. Templates were generated from blob picker performed on the p*Ph*PINK1–ADP dataset and used to pick particles from all datasets using template picker. Particles were extracted and binned (downsampled) 2×, and 2D classification was performed. Classes with any PINK1-like features were select as ‘good’ particles, and ab-initio reconstruction was performed to generate an initial reconstruction of the *Ph*PINK1 dodecamer. A second subset of classes containing particles of indistinct shapes were selected as ‘junk’ particles, and ab-initio reconstruction was performed to generate a volume for subsequent heterogeneous refinement. Three rounds of heterogeneous refinement were performed against the good and junk maps in *C*_1_ to remove bad particles from the dataset. Reconstruction of the dodecamer was performed using homogeneous refinement in *D*_3_. A mask of the dimer was generated by zoning the map from homogenous refinement against a model of the *Ph*PINK1 D357A dimer using ChimeraX (*17*, *43*), dilated and soft-padded. Particles were symmetry expanded in *D*_3_, re-centred and locally refined using the dimer mask. For the AMP-PNP-bound and ADP-bound *Ph*PINK1 datasets, signal subtraction was performed prior to local refinement. 3D variability analysis was performed solving for three modes and the particles clustered to give ∼100k particles per cluster. Particles corresponding to the most homogeneous and complete cluster were then locally refined to give the final dimer reconstruction.

Model building was performed in Coot (v0.9.8.7) (*44*) and refinement was performed using real-space refinement in Phenix (v1.20.1-4487) (*45*). The *Ph*PINK1 D357A dimer (PDB: 7T4N) (*17*) was used as the initial model and was docked into the density of the nucleotide-free *Ph*PINK1 dimer using UCSF ChimeraX (v1.6.1) (*43*). After a round of model building in Coot and refinement in Phenix, the model was docked in the densities of the *Ph*PINK1–AMP-PNP and p*Ph*PINK1–ADP dimers, and nucleotides and Mg^2+^ ions were fitted into the densities. Model building and refinement was then performed on all three models. Regions with disordered/ambiguous densities were not modelled. Cryo-EM data collection and refinement statistics are provided in Table 1.

### Thermal shift assays

Thermal shift assays were carried out using 4 μM *Ph*PINK1 (residues 115–575) and 5× SYPRO Orange Protein Gel Stain (Invitrogen) in 25 mM Tris (pH 8.5), 150 mM NaCl, 10 mM DTT, in the presence of 5 mM ADP (Sigma), AMP-PNP (Sigma or Roche), ATP (Sigma) or KTP (Biolog). 10 mM MgCl_2_ was included, unless indicated otherwise. Melt curves were measured on a Rotor-Gene Q (Qiagen) with a temperature ramp of 25–80 °C at 1 °C/min, and analysed using the Rotor-Gene Q Series Software (v2.3.1). Graphs were generated in GraphPad Prism (v9.5.1).

### Ubiquitin phosphorylation assays

Ubiquitin phosphorylation assays were carried using 1.5 μM *Ph*PINK1 (residues 115–575) and 15 μM ubiquitin in 25 mM Tris (pH 7.4), 150 mM NaCl, 10 mM MgCl_2_, 1 mM DTT. Reactions were initiated by the addition of 1 mM ATP (Sigma) or KTP (Biolog) and incubated at 22 °C for 2 h or as indicated. Reactions were quenched in SDS sample buffer (66 mM Tris (pH 6.8), 2% (w/v) SDS, 10% (v/v) glycerol, 0.005% (w/v) bromophenol blue), and samples were run on reducing 17.5% Phos-tag gels (containing 50 μM Phos-tag Acrylamide AAL-107 (Wako) and 100 μM MnCl_2_) and NuPAGE 4–12% Bis-Tris gels (Invitrogen). All gels were stained with InstantBlue Coomassie Protein Stain (Abcam).

### Cell culture and constructs

HeLa *PINK1*^−/−^ cells were a gift from Michael Lazarou (WEHI). All HeLa cell lines were cultured at 37 °C, 5% CO_2_, in DMEM supplemented with 10% (v/v) foetal bovine serum (Bovogen Biologicals) and penicillin–streptomycin. Cells were routinely checked for mycoplasma contamination using the MycoAlert Mycoplasma Detection Kit (Lonza). For transient expression, DNA encoding *Hs*PINK1 was inserted into the BamHI site of pcDNA5/FRT/TO CMVd3 vector (pcDNA5^d3^, see (*17*)) using the In-Fusion HD Cloning Kit (Takara). For stable expression, the *Hs*PINK1 sequence was inserted in between the BamHI and NheI sites of the pFU MCS SV40 Puro lentiviral vector (pFUP). Mutagenesis was performed using the Q5 Site-Directed Mutagenesis Kit (NEB).

### Generation of stable cell lines

HeLa *PINK1*^−/−^ cells stably expressing YFP–Parkin and *Hs*PINK1 were generated by sequential introduction of YFP–Parkin followed by *Hs*PINK1 into HeLa *PINK1*^−/−^ cells. YFP–Parkin was introduced using retroviral transduction with the pBMN-YFP-Parkin plasmid (gift from R. Youle; Addgene plasmid, 59416), followed by fluorescence sorting. *Hs*PINK1 WT and the M318G mutant were introduced using lentiviral transduction with pFUP-*Hs*PINK1 plasmids, followed by selection with puromycin.

### Transient transfection and Western blotting

Cells for transient transfection were seeded in 6-well plates 24–48 h prior to transfection. Transient transfections were performed with 1.5 μg pcDNA5^d3^-*Hs*PINK1 plasmids using Lipofectamine 3000 Transfection Reagent (Invitrogen), and transfected cells were allowed to grow for 24 h before harvesting. Cell lines stably expressing *Hs*PINK1 were seeded in 6-well plates 48 h before harvesting. To depolarise mitochondria to induce PINK1 stabilisation, cells were treated with 10 μM oligomycin and 4 μM antimycin A (OA) for the indicated times. Adenine and kinetin treatments were performed at 200 μM for the indicated times, and MG132 treatments were performed at 10 μM for 2 h. Cell lysates were prepared directly in SDS sample buffer, then separated on reducing NuPAGE 4–12% Bis-Tris gels (Invitrogen) or reducing 7.5% Phos-tag gels (containing 50 μM Phos-tag Acrylamide AAL-107 (Wako) and 100 μM MnCl_2_). Phos-tag gels were washed 3×10 min in 10 mM EDTA and 10 min in water prior to transfer. Protein transfer was carried out using the Trans-Blot Turbo Transfer System (Bio-Rad) onto PVDF membranes. Membranes were then blocked in 5% (w/v) skim milk powder in Tris-buffered saline containing 0.1% Tween-20 (TBS-T) and incubated with primary antibodies in TBS-T overnight at 4 °C. Membranes were washed in TBS-T, incubated in secondary antibody for ∼1 h, then washed in TBS-T prior to incubation in Clarity Western ECL Substrate (Bio-Rad) and detection using the ChemiDoc (Bio-Rad). Primary antibodies used were rabbit anti-PINK1 D8G3 (1:1,000, Cell Signaling Technology, 6946), rabbit anti-phospho-ubiquitin (Ser65) E2J6T (1:1,000, Cell Signaling Technology, 62802), rabbit anti-Tom20 FL-145 (1:1,000, Santa Cruz Biotechnology, sc-11415), mouse anti-Parkin Prk8 (1:1,000, Cell Signaling Technology, 4211). Secondary antibodies used are goat anti-rabbit HRP-conjugated (1:5,000, SouthernBiotech, 4010-05) and goat anti-mouse HRP-conjugated (1:5,000, SouthernBiotech, 1030-05). For loading controls, membranes were incubated in hFAB rhodamine anti-GAPDH (1:5,000, Bio-Rad, 12004167) overnight at 4 °C, washed in TBS-T, then detected using the ChemiDoc (Bio-Rad).

### Fractionation and ubiquitin phosphorylation assay

Ubiquitin phosphorylation assays using heavy membrane-associated *Hs*PINK1 from OA-treated HeLa *PINK1*^−/−^ cells transfected with *Hs*PINK1 WT, M318A and M318G was performed as described previously (*17*). 1×10^6^ HeLa *PINK1*^−/−^ cells were seeded in 10-cm dishes. After 48 h, cells were transfected with 5 µg pcDNA5^d3^-*Hs*PINK1 WT, M318A or M318G using Lipofectamine 3000 Transfection Reagent (Invitrogen). 24 h after transfection, *Hs*PINK1 was stabilised by OA treatment for 2 h. Cells were harvested by scraping in cold PBS and pelleted by centrifugation at 200 g for 5 min at 4 °C. Cell pellets were permeabilised by incubating for 20 min at 4 °C in 1 mL fractionation buffer (20 mM HEPES (pH 7.4), 250 mM sucrose, 50 mM KCl, 2.5 mM MgCl_2_) supplemented with 0.025% (w/v) digitonin, 1× cOmplete Protease Inhibitor Cocktail (Roche) and 1× PhosSTOP (Roche). Heavy membrane fractions were pelleted by centrifugation at 14,000 g for 5 min at 4 °C, washed once with 1 mL fractionation buffer, then resuspended in 100 µL fractionation buffer. 15 µM ubiquitin was added, and membranes were divided into 50 µL aliquots. The reaction was initiated with 1 mM ATP (Sigma-Aldrich), KTP (Biolog), or an equivalent volume of water, and incubated at 30 °C for 60 min or the indicated times with gentle agitation. Heavy membranes were pelleted by centrifugation at 14,000 g for 5 min at 4 °C, and samples for the ubiquitin containing supernatant and the *Hs*PINK1 containing heavy membrane pellet were prepared in SDS sample buffer. Western blotting was performed as described above.

### Immunofluorescence assay

HeLa *PINK1*^−/−^ cells stably expressing YFP–Parkin and *Hs*PINK1 were seeded on HistoGrip-coated coverslips 48 h prior to treatment with OA and/or kinetin for the indicated times. Cells were then fixed with 4% (w/v) paraformaldehyde (PFA) in 0.1 M phosphate buffer on a rocker for 10 min, rinsed three times with PBS and permeabilised with 0.1% (v/v) Triton X-100 in PBS for 10 min. After that, samples were blocked with 3% (v/v) goat serum in 0.1% (v/v) Triton X-100/PBS for 15 min and incubated with anti-GFP (ThermoFisher, A10262) and anti-mitochondrial HSP60 (Abcam, ab128567) antibodies for 90 min. Following three washes with PBS and subsequent 1 h incubation with Alexa Fluor 488 goat anti-chicken IgG (ThermoFisher, A32931) and Alexa Fluor 647 goat anti-mouse IgG (ThermoFisher, A21235), the coverslips were washed three times with PBS and mounted with a Tris-buffered DABCO-glycerol mounting medium onto glass slides. Imaging of the coverslips was done with an inverted Leica SP8 confocal laser scanning microscope under 63×/1.40 NA objective (Oil immersion, HC PLAPO, CS2; Leica microsystems). Details on the staining procedure are available at (*46*).

## Author Contributions

ZYG performed all experiments except as listed hereafter and analysed data. SC and TNN performed mitophagy imaging experiments. AL and NSK collected and processed cryo-EM data. ML, GD and DK supervised the work and obtained funding. ZYG and DK wrote the manuscript with input from all authors.

## Acknowledgements

We thank Jeffrey Babon and Isabelle Lucet for helpful advice. We acknowledge use of the facilities at the Ian Holmes Imaging Centre, Bio21 Institute. Funding in the author’s labs is provided by the National Health and Medical Research Council (GNT1178122 to DK, GNT2004446 to GD, GNT1106471 to ML), the Australian Research Council (DP200100347 to ML), the Michael J Fox Foundation (DK, GD), the Bodhi Education Fund (GD) and an Australian Government Research Training Program Fellowship (to ZYG).

## Data and materials availability

All data, code, and materials used in the analyses are available upon reasonable request from the corresponding author. Structures have been submitted to relevant repositories (pdb, EMDB). All data are available in the main text or the supplementary materials.

---

**Supplementary Figure 1.**
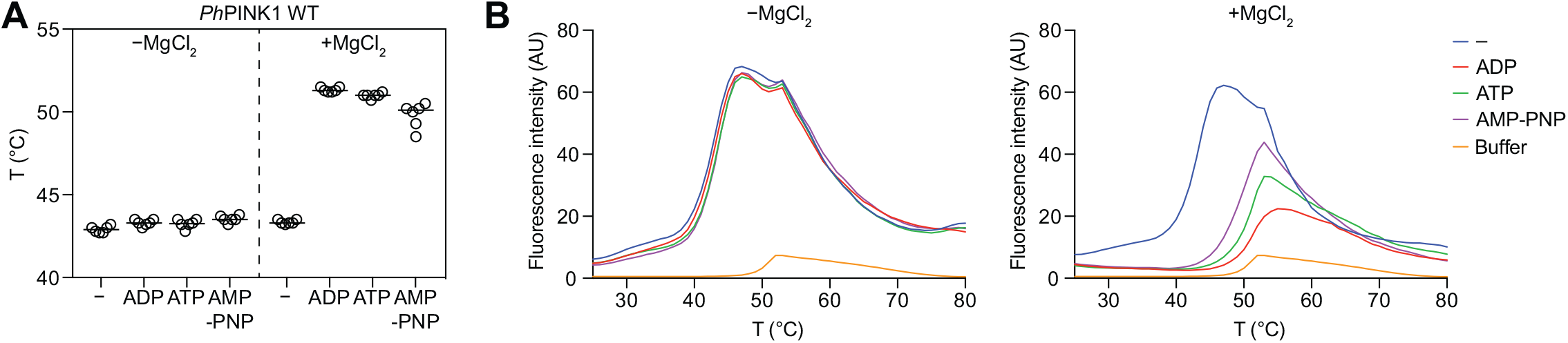
Thermal stability of WT *Ph*PINK1 in the presence of nucleotides and Mg^2+^. **(A)** Melting temperatures of WT *Ph*PINK1 (residues 115–575, monomeric, autophosphorylated) in the presence of ADP, ATP or AMP-PNP, with/without MgCl_2_. All nucleotides bind and stabilise *Ph*PINK1 in the presence of MgCl_2_. Experiment was performed three times in technical duplicates. **(B)** Representative thermal melt curves for the data in **A**. Note that buffer alone (no protein, nucleotide or MgCl_2_) produces a small melt curve that overlaps with but is unlikely to significantly impact the melt curves of nucleotide bound *Ph*PINK1. The melt curves for *Ph*PINK1 in the absence of MgCl_2_ (left) and in the presence of MgCl_2_ (right) were generated in the same experiment, and are separated for clarity; the buffer curve shown is the same between the two graphs.

**Supplementary Figure 2.**
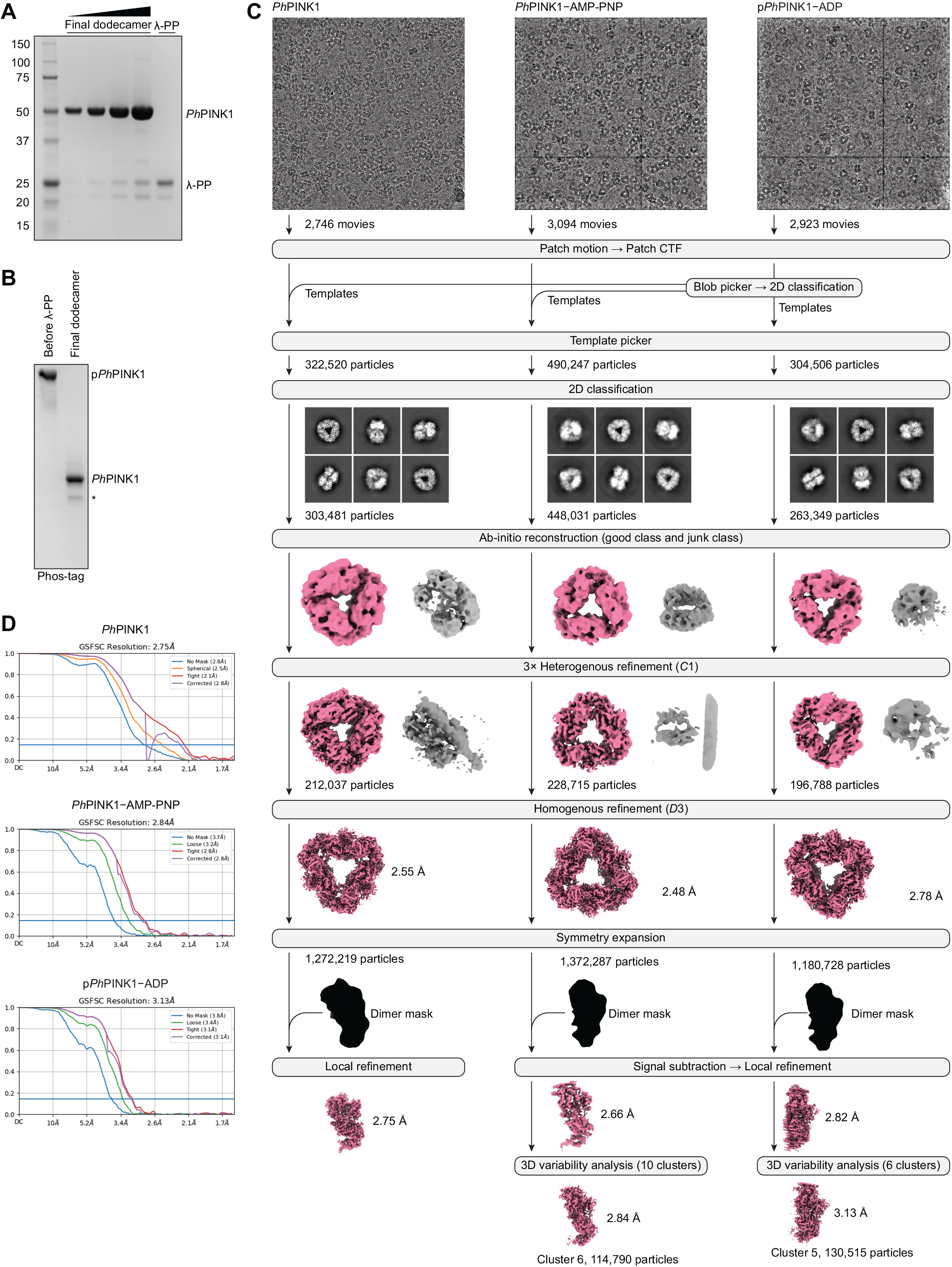
Cryo-EM analysis of nucleotide-bound *Ph*PINK1. **(A)** The final purified dephosphorylated WT *Ph*PINK1 dodecamer run at increasing concentrations on SDS-PAGE, revealing a minor contaminating species at approximately 25 kDa, likely to be λ-PP. **(B)** Phos-tag analysis of *Ph*PINK1 before dephosphorylation by λ-PP, and after purification of the final *Ph*PINK1 dodecamer. Note, the faster migrating band (labelled with an asterisk) is likely to be fully dephosphorylated *Ph*PINK1, while the major band is presumed to be *Ph*PINK1 pThr305 given that density for pThr305 can be seen in subsequent cryo-EM reconstructions (Supplementary Figure 3C). Dephosphorylation of pThr305 is likely prevented by the adjacent Pro306 (*18*, *47*). Phosphorylation of Thr305 is a by-product of non-specific *Ph*PINK1 autophosphorylation during expression in bacteria (*18*). **(C)** Cryo-EM processing pipeline for the nucleotide-free, AMP-PNP-bound and ADP-bound *Ph*PINK1 dimers. Representative micrographs and select 2D classes are shown. **(D)** Gold standard Fourier shell correlation (GSFSC) curves for each of the final maps in **C**.

**Supplementary Figure 3.**
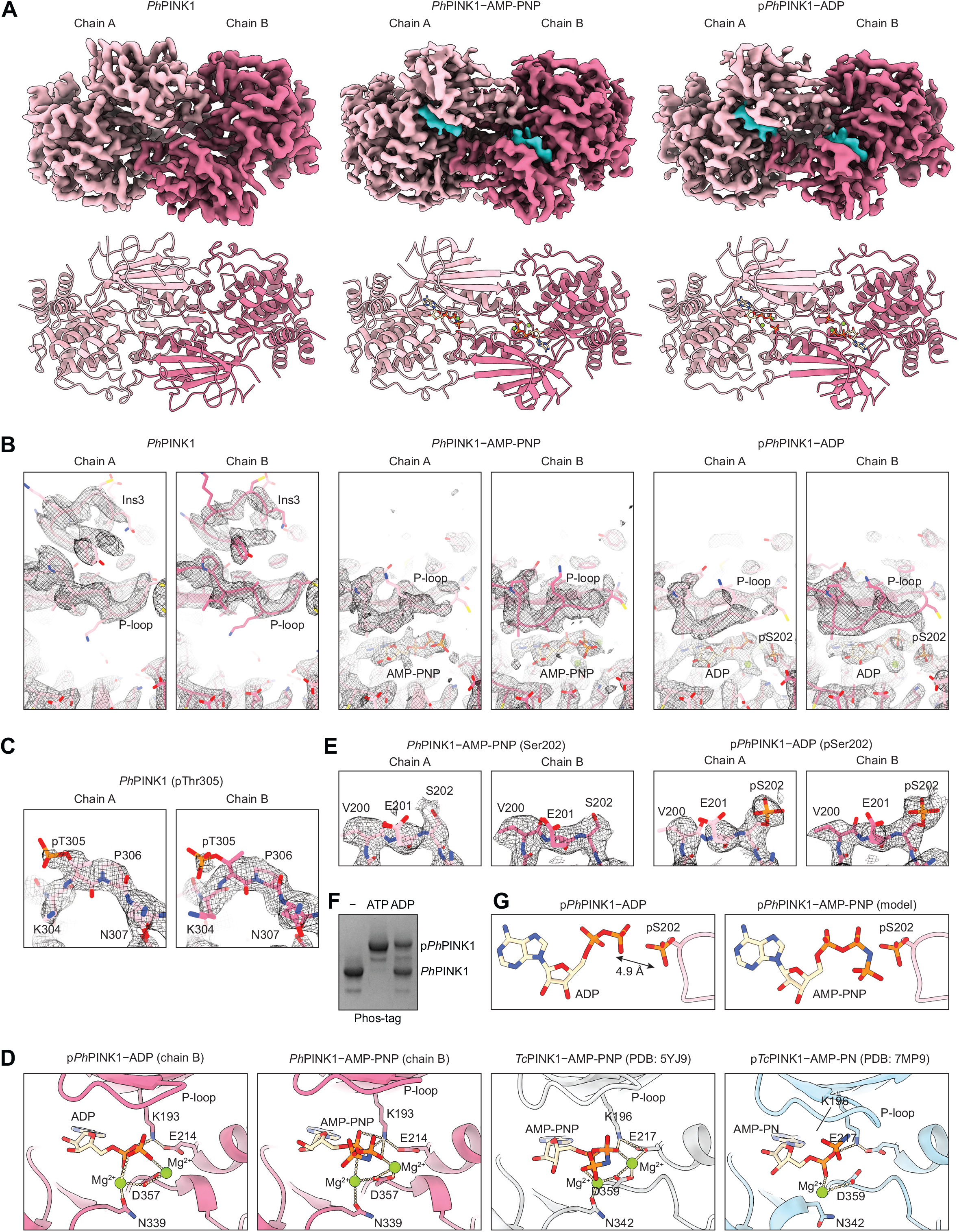
Cryo-EM structure of the nucleotide-bound *Ph*PINK1 dimers. **(A)** Cryo-EM densities (*top*) and cartoon model (*bottom*) of the nucleotide-free, AMP-PNP-bound and ADP-bound *Ph*PINK1 dimers. Each *Ph*PINK1 monomer is coloured in a different shade of pink, and density for the nucleotide and Mg^2+^ ions is coloured in cyan. Densities for nucleotide-free, AMP-PNP-bound and ADP-bound *Ph*PINK1 are contoured to levels 0.35, 0.12 and 0.12, respectively. **(B)** Zoomed view of the ATP binding site, P-loop and insertion-3 of chains A and B of each *Ph*PINK1 dimer. Residue side chains and nucleotides are shown as sticks, and Mg^2+^ as green spheres. Corresponding densities are displayed as a mesh. **(C)** Density for phosphorylated Thr305 can be seen in each chain of the nucleotide-free *Ph*PINK1 dimer, consistent with Phos-tag analysis of the purified dephosphorylated *Ph*PINK1 dodecamer that was used for cryo-EM (see Supplementary Figure 2B and legend) and previous observations (*18*). **(D)** Comparison between the binding *Ph*PINK1 to AMP-PNP and ADP (only chain B is shown) and the binding of *Tc*PINK1 to AMP-PNP (PDB: 5YJ9, (*21*)) and AMP-PN (PDB: 7MP9, (*19*)). Polar interactions are shown as dotted lines. *Ph*PINK1 and *Tc*PINK1 bind nucleotides via similar interactions. **(E)** Density for phosphorylated Ser202 can be seen for both chains of p*Ph*PINK1– ADP, but not *Ph*PINK1–AMP-PNP. **(F)** The dephosphorylated *Ph*PINK1 dodecamer was incubated with 10 mM ADP (or ATP as a positive control) and 10 mM MgCl_2_ for 10 min on ice. The ADP stock used in the experiment was the same stock used for cryo-EM analysis. Phos-tag analysis revealed that ADP incubation leads to *Ph*PINK1 autophosphorylation, indicating that the ADP stock is contaminated with ATP and explaining the density for pSer202 seen in cryo-EM reconstructions (see **E**). **(G)** *Left*, the position of ADP (chain B) relative to pSer202 (chain A) in the p*Ph*PINK1–ADP structure. *Right*, AMP-PNP is modelled using pSer202 from the p*Ph*PINK1–ADP.

**Supplementary Figure 4.**
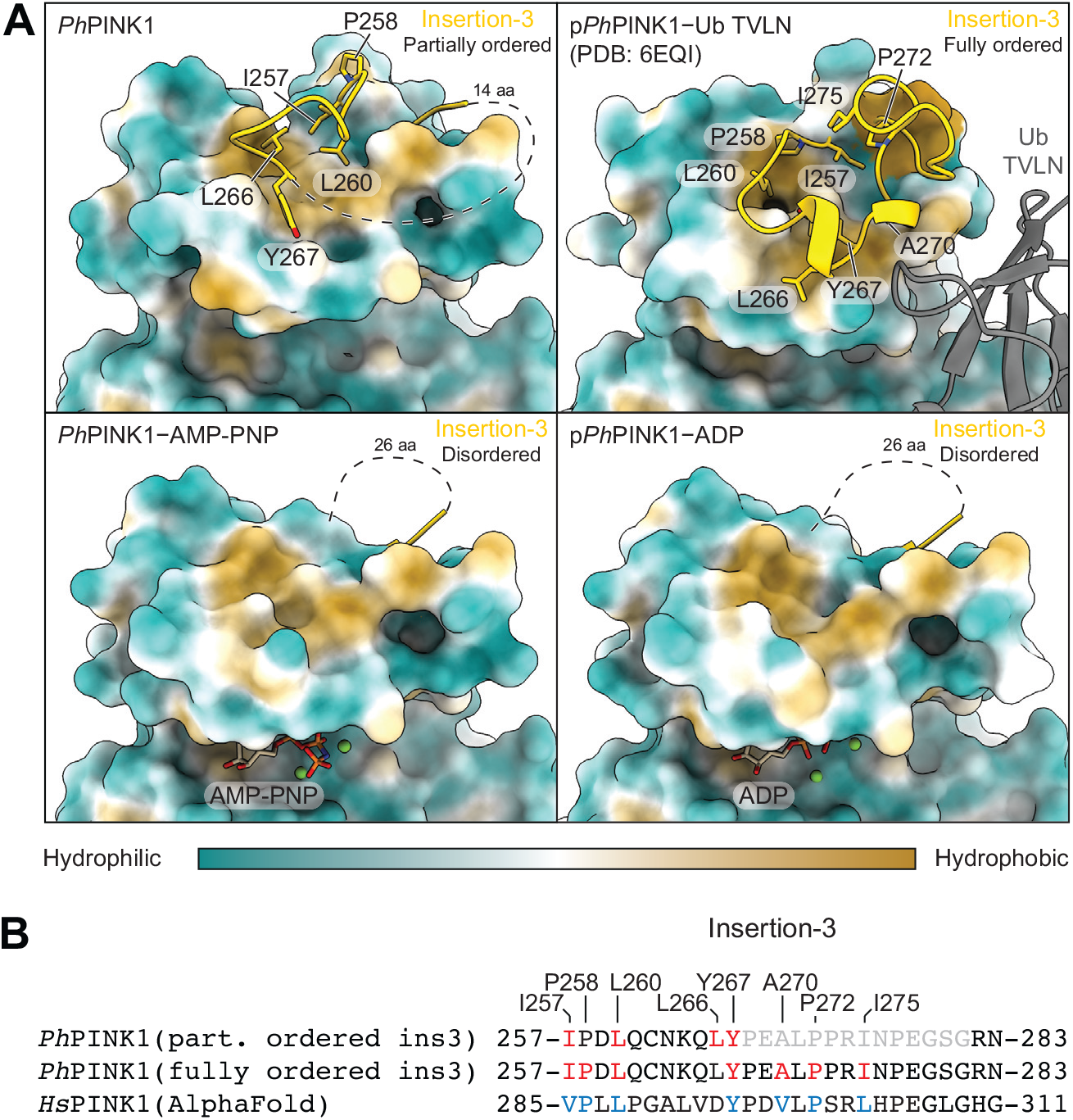
The hydrophobic interface between insertion-3 and the N-lobe. **(A)** Hydrophobicity of the N-lobe of *Ph*PINK1 in the nucleotide-free, AMP-PNP-bound and ADP-bound states (only chain B is shown), and the phosphorylated and ubiquitin-bound *Ph*PINK1 complex (PDB: 6EQI, (*18*)). Insertion-3, shown in cartoon representation, shields a hydrophobic patch in the N-lobe. Hydrophobic residues are shown as sticks. aa, amino acids. **(B)** Sequence alignment of insertion-3 from *Ph*PINK1 and *Hs*PINK1. *Ph*PINK1 residues involved in the hydrophobic insertion-3–N-lobe interaction are highlighted in red. *Hs*PINK1 residues predicted by AlphaFold (*48*, *49*) to be involved in the hydrophobic insertion-3–N-lobe interaction are highlighted in blue. Residues in light grey are disordered in the structure. ins3, insertion-3.

**Supplementary Figure 5.**
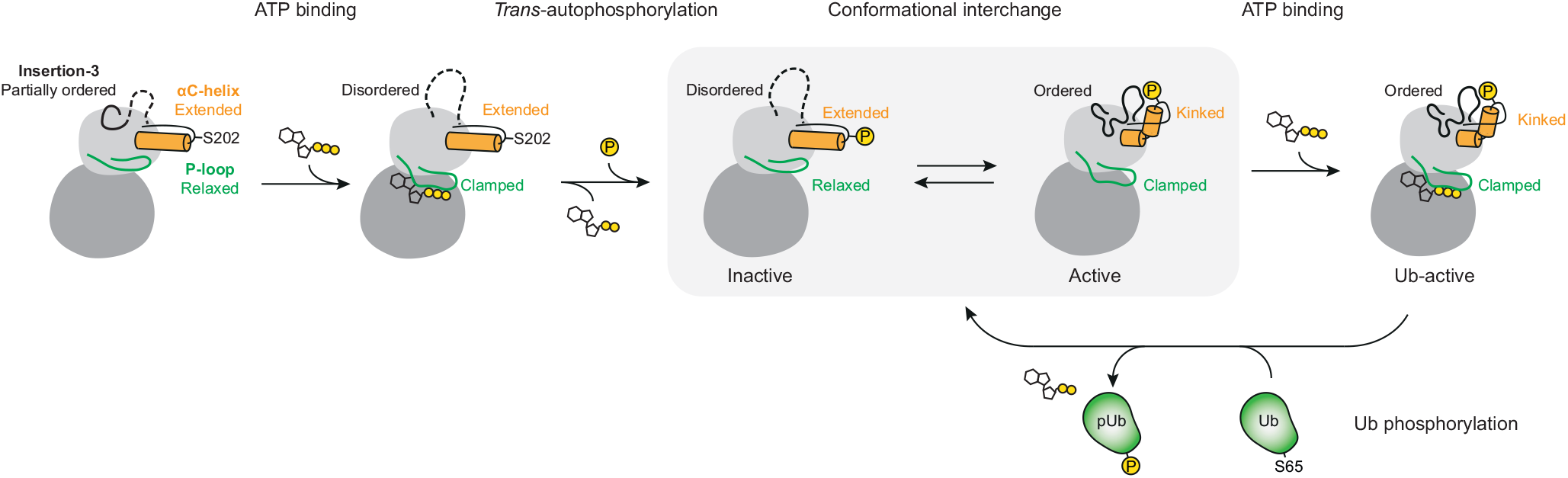
Model of how ATP binding contributes to PINK1 activation and ubiquitin phosphorylation. Unphosphorylated PINK1 dimerises and utilises ATP to *trans*-autophosphorylate at Ser202. The resulting phosphorylated PINK1 can adopt two conformations, an inactive conformation that cannot bind ubiquitin (relaxed P-loop, extended αC-helix, disordered insertion-3) and an active conformation that can bind ubiquitin (clamped P-loop, kinked αC-helix, ordered insertion-3). Binding of ATP to phosphorylated PINK1 stabilises the active conformation via its interaction with the clamped P-loop, enabling efficient ubiquitin recognition and phosphorylation by PINK1.

**Supplementary Figure 6.**
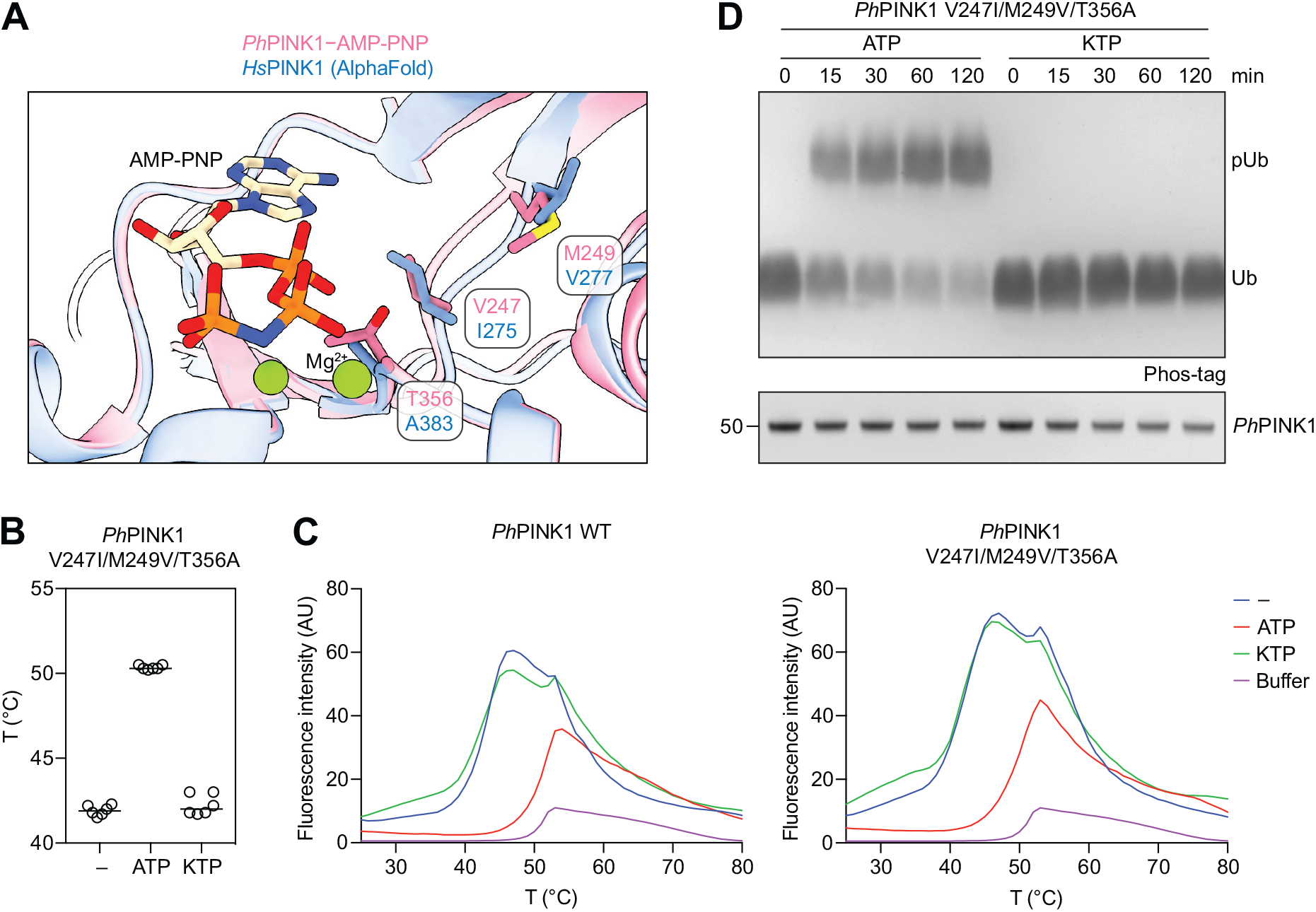
A humanised mutant of *Ph*PINK1 does not bind KTP. **(A)** Superimposed ATP binding sites of *Ph*PINK1–AMP-PNP and *Hs*PINK1 (AlphaFold; (*48*, *49*)). The N-lobe β1-, β2- and β3-strands are hidden for clarity. Three AMP-PNP-proximal residues in *Ph*PINK1 (Val247, Met249, Thr356) were mutated into their equivalent *Hs*PINK1 counterparts to generate a humanised version of *Ph*PINK1 for the experiments in **B–D**. **(B)** Melting temperatures of *Ph*PINK1 V247I/M249V/T356A in the presence of ATP or KTP. No increase in melting temperature is observed for KTP, indicating an absence of interaction. Experiment was performed three times in duplicate. **(C)** Representative thermal melt curves for the data in **B** and Figure 3B. The buffer curve was generated in the absence of protein and nucleotide. The melt curves for WT *Ph*PINK1 and *Ph*PINK1 V247I/M249V/T356A were separated into different graphs for clarity, and the buffer curve shown is the same between the two graphs. **(D)** Ubiquitin phosphorylation assay using *Ph*PINK1 V247I/M249V/T356A in the presence of ATP or KTP, analysed on a Phos-tag gel. *Ph*PINK1 activity was not detected when KTP was supplied. Experiment was performed in triplicate.\

**Supplementary Figure 7.**
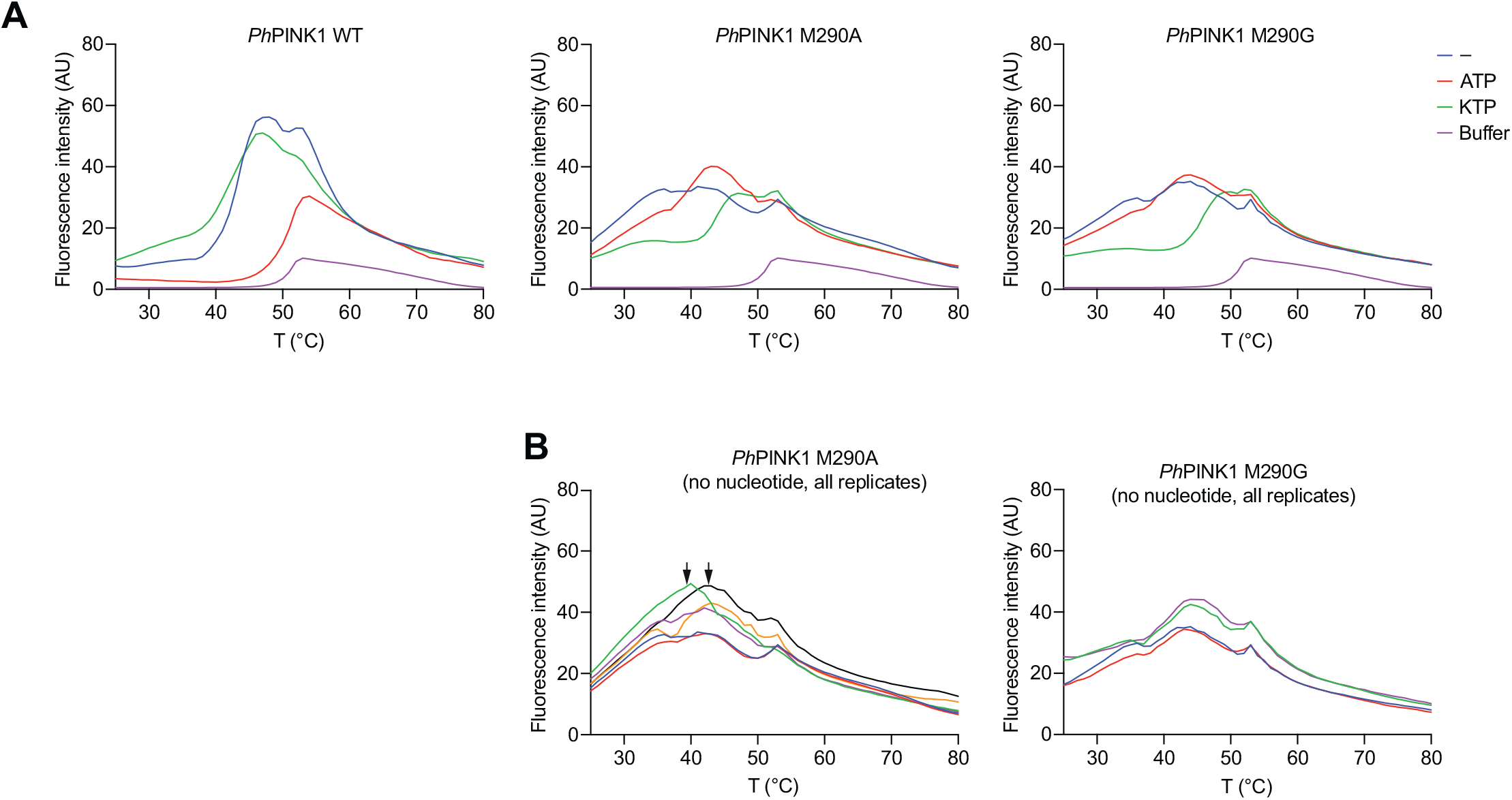
Instability of *Ph*PINK1 M290A and M290G. **(A)** Representative thermal melt curves for experiment in Figure 3F. *Ph*PINK1 M290A and M290G (*middle* and *right*, respectively) in the absence of nucleotide (blue curves) or in the presence of ATP (red curves) show relatively indistinct melt curves, indicating low stability of either mutant. Addition of KTP resulted in better defined melt curves (green curves), consistent with nucleotide binding increasing the stability of the M290A and M290G mutants. The experiment was repeated a total of two (WT, M290G) or three (M190A) times in technical duplicates. Melting temperatures could only be calculated for a subset of replicates (see **B**). Melt curves for all three *Ph*PINK1 variants were separated for clarity; the buffer curve shown is the same between the three graphs. **(B)** All replicate melt curves for M290A and M290G in the absence of nucleotide are shown. While melting temperatures could not be determined for two of the M290A curves (arrows), the other four curves displayed a small inflection at ∼40 °C that was sufficient for melting temperature calculation.

**Supplementary Figure 8.**
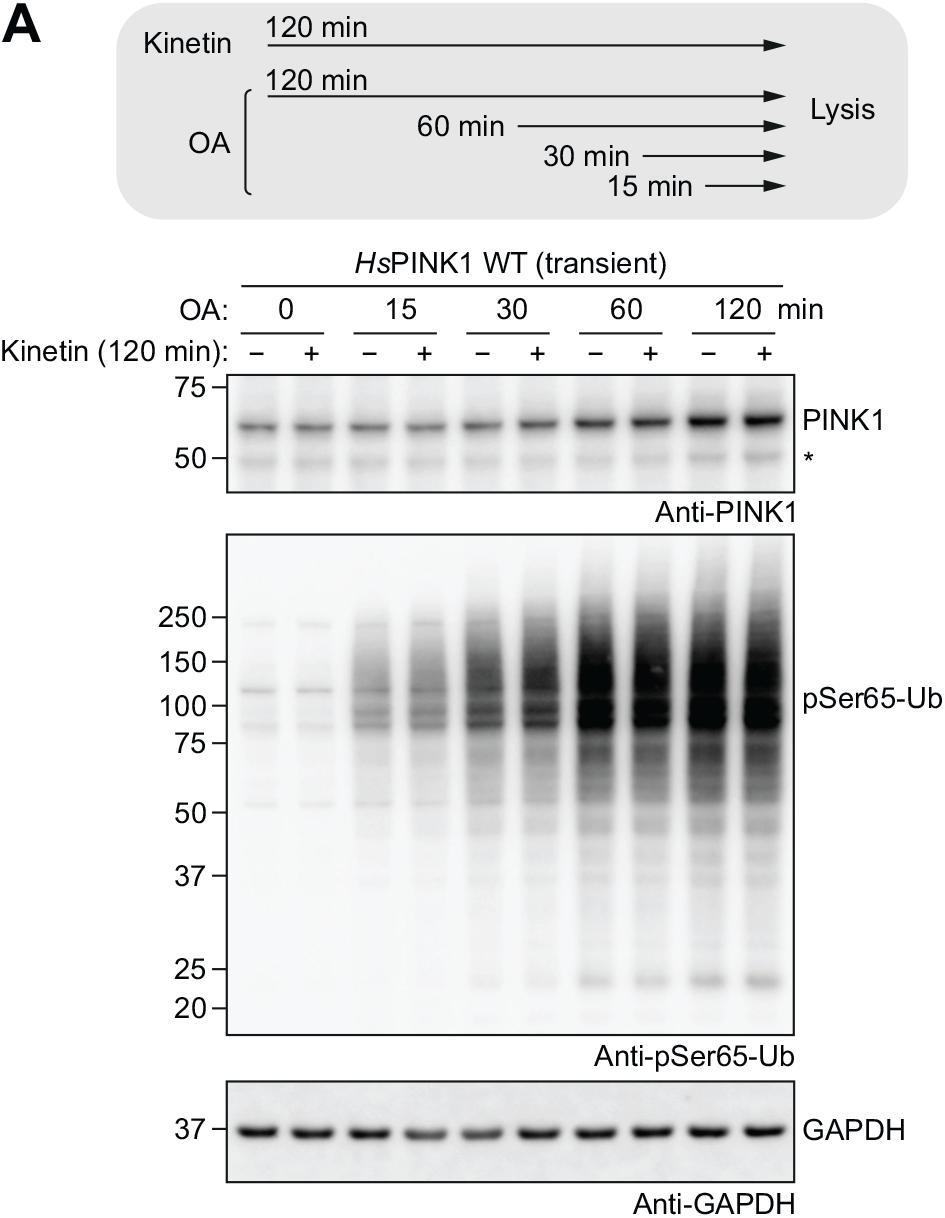
Kinetin does not appear to affect the ubiquitin kinase activity of *Hs*PINK1 WT. HeLa *PINK1*^−/−^ YFP–Parkin cells transiently expressing WT *Hs*PINK1 were co-treated with OA and kinetin according to the schematic (top panel). Immunoblotting shows that kinetin has no effect on WT *Hs*PINK1-mediated ubiquitin phosphorylation. The asterisk indicates the 52-kDa PARL-cleaved PINK1. Experiment was performed in triplicate.

**Supplementary Figure 9.**
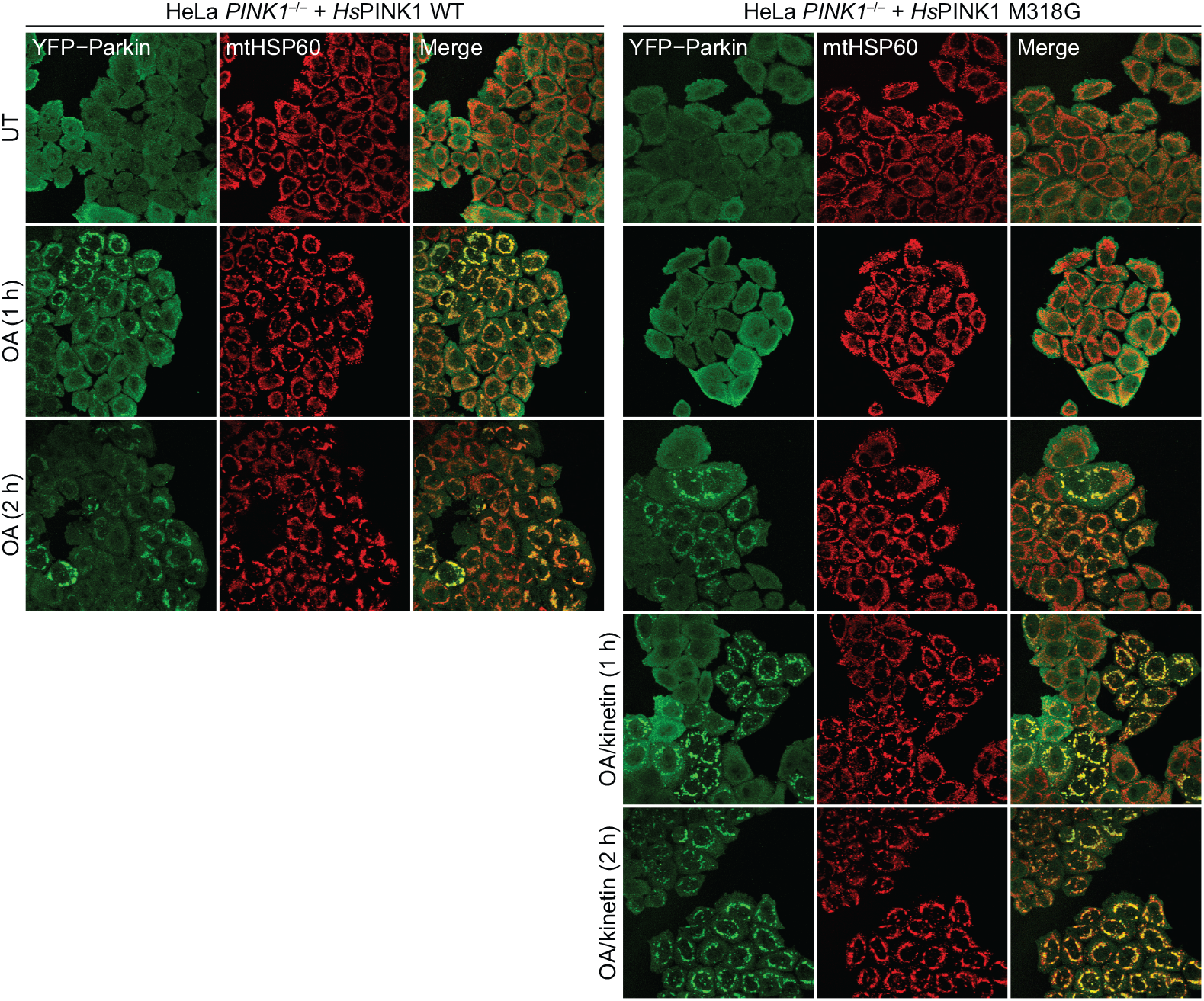
Repeat of Figure 5 but showing the 2 h OA/kinetin treatment timepoints. At 2 h, OA treatment alone induced YFP–Parkin translocation in *Hs*PINK1 M318G-expressing cells. In this repeat, cells were additionally immunostained for PINK1, but fluorescence signal consistent with PINK1 was not detected and therefore not shown. Experiment was performed in duplicate with Figure 5.

